# Genetic Allee effects and their interaction with ecological Allee effects

**DOI:** 10.1101/061549

**Authors:** Meike J. Wittmann, Hanna Stuis, Dirk Metzler

## Abstract

**Summary:**

1. It is now widely accepted that genetic processes such as inbreeding depression and loss of genetic variation can increase the extinction risk of small populations. However, it is generally unclear whether extinction risk from genetic causes gradually increases with decreasing population size or whether there is a sharp transition around a specific threshold population size. In the ecological literature, such threshold phenomena are called “strong Allee effects” and they can arise for example from mate limitation in small populations.
2. In this study, we aim to a) develop a meaningful notion of a “strong genetic Allee effect”, b) explore whether and under what conditions such an effect can arise from inbreeding depression due to recessive deleterious mutations, and c) quantify the interaction of potential genetic Allee effects with the well-known mate-finding Allee effect.
3. We define a strong genetic Allee effect as a genetic process that causes a population’s survival probability to be a sigmoid function of its initial size. The inflection point of this function defines the critical population size. To characterize survival-probability curves, we develop and analyze simple stochastic models for the ecology and genetics of small populations.
4. Our results indicate that inbreeding depression can indeed cause a strong genetic Allee effect, but only if individuals carry sufficiently many deleterious mutations (lethal equivalents) on average and if these mutations are spread across sufficiently many loci. Populations suffering from a genetic Allee effect often first grow, then decline as inbreeding depression sets in, and then potentially recover as deleterious mutations are purged. Critical population sizes of ecological and genetic Allee effects appear to be often additive, but even superadditive interactions are possible.
5. Many published estimates for the number of lethal equivalents in birds and mammals fall in the parameter range where strong genetic Allee effects are expected. Unfortunately, extinction risk due to genetic Allee effects can easily be underestimated as populations with genetic problems often grow initially, but then crash later. Also interactions between ecological and genetic Allee effects can be strong and should not be neglected when assessing the viability of endangered or introduced populations.

## Introduction

Life can be challenging for the members of a small or sparse population (Fauvergue *et al*. 2012). While many plants suffer from pollen limitation, animals may struggle to find mating partners (Gascoigne *et al*. 2009) or cooperators for hunting, defense, and rearing offspring (Courchamp *et al*. 1999). Individuals in small populations may also be more likely to be killed by a predator (Kramer & Drake 2010; McLellan *et al*. 2010). Moreover, small populations are at a high risk of going extinct due to random fluctuations in the number of birth and death events, sex ratio, and environmental conditions (Fauvergue *et al*. 2012).

In addition to these ecological problems, small populations can also experience a number of genetic problems, including inbreeding depression, mutation accumulation, and the loss of genetic variation and evolutionary potential (Frankham 2005). While some small populations may only suffer from ecological problems, and some only from genetic problems, there may be many small populations suffering from both at the same time. For example, in metapopulations of the Glanville fritillary butterfly, *Melitaea cinxia*, small populations have reduced mating success and a higher emigration rate (Kuussaari *et al*. 1998). In addition, there is evidence for inbreeding depression (Saccheri *et al*. 1998). Similarly, the rare Marsh Gentian, *Gentiana pneumonanthe*, exhibits both pollen limitation and inbreeding depression (Oostermeijer 2000).

With the exception of demographic and environmental stochasticity, the above-mentioned ecological problems can cause a “demographic Allee effect”, defined as a reduced average percapita growth rate in small populations (Stephens *et al*. 1999). The “demographic” indicates that population growth rate is affected and not just one fitness component. But since this study is only concerned with demographic Allee effects, we will simply refer to them as Allee effects in the following.

There are two types of Allee effect with qualitatively different population dynamics: weak and strong Allee effects (Wang & Kot 2001; Taylor & Hastings 2005). Under a weak Allee effect, the per-capita growth rate is reduced but still positive in very small populations. Thus populations of any size can survive. Under a strong Allee effect, the per-capita growth rate is negative for populations below a certain critical population size. In a deterministic world, populations starting out below this critical population size are doomed to extinction.

For a more realistic stochastic setting, Dennis (1989, 2002) demonstrated that, independently of the form of stochasticity, a strong Allee effect leads to a characteristically sigmoid (S-shaped) graph of success probability, i.e. the probability to escape extinction, as a function of founder population size. Success probability first increases very slowly with increasing founder population size, then increasingly rapidly with an inflection point at the critical population size of the corresponding deterministic model, and then levels of as it is approaching one. Neither a weak Allee effect nor demographic or environmental stochasticity alone produce such a signature. In these latter cases, success probability first increases rapidly with increasing initial population size and then levels off (Dennis 2002). Even in a stochastic setting, it is of course highly relevant for the management of an endangered or invasive species whether extinction risk gradually increases with decreasing population size or whether there is a specific threshold population size around which there is a sharp increase in extinction risk.

Although the importance of genetic factors in species extinctions has been questioned in the past, most prominently by Lande (1988), it is now widely accepted that genetic processes acting in small populations do contribute to driving small populations to extinction (Frankham *et al*. 2002; Spielman *et al*. 2004; O’Grady *et al*. 2006). A number of times, such genetic processes have therefore been referred to as “genetic Allee effects” (e.g., Fischer *et al*. 2000; Willi *et al*. 2005; Courchamp *et al*. 2008; Fauvergue *et al*. 2012). However, there are at least two reasons to be cautious with such a designation. First, as illustrated by the case of demographic or environmental stochasticity, not everything that leads to increased extinction risk in small populations is also an Allee effect, or even a strong Allee effect with a critical population size. Second, the original definition of the Allee effect is based on the per-capita growth rate in small populations, and it is unclear how to apply this definition to situations with genetic change where per-capita growth rate is not a function of population size alone. Two populations of the same size (or even the same population at different time points) generally have different growth rates depending on their genetic composition, for example their level of inbreeding.

Here we employ the results by Dennis (1989, 2002) to make progress on defining and detecting genetic Allee effects. Our approach specifically targets strong Allee effects, i.e. those most relevant to the survival of a population. We define a “strong genetic Allee” effect as a genetic process that produces an inflection point (S-shape) in the graph of persistence probability as a function of initial population size. With this definition, it is not a problem that the genetic composition and growth rate of a population change over time, but we need to keep in mind that all results will be relative to the genetic composition of the source population.

We focus on one important candidate mechanism for genetic Allee effects: inbreeding depression due to recessive deleterious mutations (Charlesworth & Willis 2009). In large outcrossing populations, such mutations do not cause many deaths because almost all individuals are heterozygous. However, when such a population is suddenly reduced to small size, mating between close relatives becomes more likely in the subsequent generations and thus the recessive mutations are more often found in homozygous state. This reduces individual viability and can lead to a reduction in population growth rate. Indeed, laboratory experiments on mice and flies (Frankham 1995), field studies on a butterfly metapopulation (Saccheri *et al*. 1998), as well as previous modeling results (Tanaka 1997, 1998, 2000; Robert 2011) suggest that inbreeding depression can indeed contribute to driving populations to extinction. It is still unclear, however, whether the consequences of inbreeding depression can be described as a strong Allee effect with a critical population size. One complication is that recessive deleterious mutations are increasingly exposed to selection in shrinking populations. Thus, their frequency will decrease over time and, if the population does not go extinct, viability will eventually rebound. The population growth rate may then exceed that of larger populations that could not “purge” their deleterious mutations (Wang *et al*. 1999).

We consider populations that suddenly find themselves at small size or low density, either as a small sample from a larger population introduced into a new habitat or as a long-term resident that just experienced a catastrophic crash in population size or a fragmentation of its range. Using a stochastic model for the population dynamics and genetics of such a population, we first determine under what conditions recessive deleterious mutations cause a strong genetic Allee effect, i.e. a sigmoid success-probability curve. Second, we characterize the interaction of ecological and genetic Allee effects acting in the same population.

## Methods

### Model

We constructed a stochastic individual-based simulation model for the population dynamics of a small founder population, which may experience mate-finding difficulties and reduced viability due to recessive deleterious mutations (see Table 1 for a list of variables and parameters). Our model population is diploid, there are males and females, and reproduction is obligately sexual. We assume random mating, such that any inbreeding is due to small population size (panmictic inbreeding rather than systematic inbreeding *sensu* Glémin 2003). The simulation proceeds in discrete time steps. In each generation, events take place in the following order: mate finding, reproduction with inheritance of genetic material, and finally death of the parent generation and replacement by the offspring generation.

**Table 1:** List of model variables and parameters used in the main text

| Parameter/Variable | Explanation |
| --- | --- |
| *Population level* |  |
| $N_0$ | Size of founder population |
| $F_t$ | Number of females in generation $t$ |
| $M_t$ | Number of males in generation $t$ |
| $r$ | Encounter rate |
| $w$ | Fecundity, i.e., average number of offspring produced by a mated female |
| $b$ | Number of lethal equivalents (LE); here the average number of lethal mutations per haploid genome |
| $n$ | Number of loci with recessive lethal mutations |
| $q$ | Frequency of the recessive lethal alleles in the source population |
| *Individual level* |  |
| Sex | Male or female |
| $d_i$ | Number of lethal allele copies at locus $i$ with $i \in \{1, \dots, n\}$ |

### Properties of individuals

Individuals are characterized by their sex (male or female) and by their genotype. We assume that there are *n* unlinked loci with completely recessive lethal mutations, the simplest possible genetic basis for inbreeding depression. An individual’s copy number of the lethal allele at locus *i* with *i* = 1, …, *n* is denoted *d_i_* and - since individuals are diploid - takes values in {0, 1, 2}. Individuals homozygous for the respective lethal allele at one or more loci are not viable. In Appendix S3, we explore several other possible genetic architectures (see below for further explanation).

### Constructing the founder population

Each of the *N*_0_ individuals in the founder population is independently and with equal probability male or female. The key parameter for the genetic composition of the founder population is the number of “lethal equivalents” in the source population, *b*. In general, one lethal equivalent represents either one fully recessive lethal mutation per haploid genome or a set of mutations of smaller effect or larger dominance that jointly cause the same amount of inbreeding depression. Here we assume fully recessive lethals and therefore *b* is simply the average number of lethal alleles per haploid genome in the source population (Appendix S3 explores more complex genetic architectures and Appendix S3.1 explains how the number of lethal equivalents is defined in such cases). The lethal alleles are distributed evenly across loci, such that each locus has lethal allele frequency *q* = *b*/*n* (see Appendix S3 for an extended model with variation in allele frequencies across loci). For each individual and each locus independently, we draw the number of lethal allele copies from a binomial distribution with parameters *q* and 2 (because of diploidy). Because we assume that the founder individuals are drawn from the viable members of the source population, we discard individuals that are homozygous for the lethal allele at one or more loci, and repeat the random drawing of individual genotypes until the required number of viable founder individuals is reached.

### Mate finding

Independently of all other females, each female in generation *t* finds a mating partner with probability 1 − *e*^−*r·M_t_*^, where *r* is the per-capita encounter rate and *M_t_* is the number of males in generation *t*. Under the assumption that the time until the first encounter is exponentially distributed, this is the probability that a female encounters at least one male during its lifetime of one generation. We assume here that each female mates with only one male or equivalently that all her offspring is sired by the male she mated with first. For each female independently, all males have the same chance to be encountered first, i.e. they can also mate with multiple females. In Appendix S1, we explore an alternative model in which females can also mate multiple times and in Appendix S2 we consider a hermaphroditic model.

### Reproduction

Each mated pair independently produces a Poisson-distributed number of offspring, *X*, with mean *w*, the fecundity parameter. For each of the *X* offspring independently, the sex is assigned randomly (male or female with equal probability) and the genotype is constructed by randomly drawing from the two parents’ genomes according to the Mendelian rules and assuming free recombination between loci. That is, for each locus the parents pass on one of their two allele copies with equal probability. If an offspring individual receives a lethal allele copy from both mother and father at one or more loci it is discarded as not viable. Thus the number of offspring surviving to the next generation may be smaller than the initially drawn number, *X*. Since we are interested in the fate of a small population on a short time scale, new mutations should play a minor role and are neglected.

### End of the life cycle

At the end of each generation, all viable female and male offspring are counted to determine the number of females and males in the next generation, *F*_*t*+1_ and *M*_*t*+1_. Finally, the individuals in the parent generation die and the offspring individuals become the new parents for the next generation.

### Simulations

To understand the consequences of ecological and genetic problems individually, as well as their interaction, we compared four sets of simulations. In the first set (**D**), there was neither mate limitation (*r* = ∞) nor were there genetic problems due to lethal mutations (*b* = 0), i.e. all offspring were viable. Small populations could only go extinct through demographic stochasticity, for example because by chance none of the mated pairs produced any offspring or because all individuals in a generation were of the same sex. Demographic stochasticity is present in all our simulations. In the second set (**D+E**), there were additional ecological problems, i.e. mate-finding problems with encounter rate *r* < ∞, but no genetic problems (*b* = 0). In the third set (**D+G**), there were additional genetic problems (*b* > 0), but no mate-finding difficulties (*r* = ∞). Finally, in the full model (**D+E+G**), populations could experience both genetic problems (*b* > 0) and mate-finding problems (*r* < ∞).

We ran these four sets of simulations for various parameter combinations. In each case, we simulated 1,000 colonization attempts each for population sizes between 5 and 995 in steps of five. A simulation run was stopped once the population either reached a size of 1,000 individuals, in which case we call the population successful, or went extinct. The proportion of successful populations will be a good proxy for long-term survival probability if we can assume that populations reaching 1,000 individuals will keep growing and not succumb to inbreeding depression at a later stage. By repeating simulations with a target size of 2,000 individuals, we confirmed that this assumption is indeed reasonable.

To explore ranges of parameter values that are most meaningful for natural populations, we considered empirical estimates for the number of lethal equivalents, *b*. These estimates are generally based on the slope of the relationship between inbreeding coefficient and the logarithm of viability (Morton *et al*. 1956, see also Appendix S3). In a meta-analysis by O’Grady *et al*. (2006), the average value of *b* for birds and mammals was estimated to be around six. However, there appears to be wide variation among species (see O’Grady *et al*. 2006, Table 1 and references therein), and also among environments, with higher values of *b* in more stressful environments (Liao & Reed 2009). By considering values of *b* up to 12, we should represent most of the range observed in natural populations.

Unfortunately, the genetic architecture underlying a certain number of lethal equivalents, i.e. the number of contributing loci and their distribution of selection and dominance coefficients, is generally unknown and it is unclear to what extent it is important. Miklos & Rubin (1996) estimated the number of loci with lethal mutations in the genome to be on the order of thousands for a number of animal species, and on the order of hundreds for *Arabidopsis thaliana*. Here we focus on fully recessive lethal mutations and compare two values for the underlying number of loci, *n*, 100 and 1,000. In Appendix S3, we explore three alternative genetic architectures underlying a given number of lethal equivalents. First, we assumed a set of lethal mutations that all have dominance coefficient *h* = 0.05. In natural populations, however, inbreeding depression will be due to the combined effects of many loci with different parameters. Second, we therefore considered a distribution of selection and dominance coefficients estimated for yeast (Agrawal & Whitlock 2011), the only species, to our knowledge, for which such detailed information is currently available. Third, we assumed that half of the lethal equivalents are due lethal mutations and the other half due to mutations of smaller effect, as has been suggested for *Drosophila* (Mukai *et al*. 1972). In the latter two cases we also took into account that mutations can have different equilibrium frequencies in the source population, depending partly on their parameters and partly on chance effects. We hypothesized that different genetic architectures with the same number of lethal equivalents will exhibit a similar degree of inbreeding depression initially, but that they could differ substantially in how readily deleterious mutations can be purged by small populations. For example, small-effect mutations might be more difficult to purge than lethal mutations (Wang *et al*. 1999).

### Analysis

For each set of 1,000 replicate simulations, we computed the proportion of successful populations, *P*. Since the initial genetic configuration is drawn independently for each replicate population, *P* represents an average over possible configurations of the founder population, given the number of lethal equivalents, *b*, in the source population. To visually check for the signature a strong Allee effect, we plotted *P* as a function of founder population size, *N*_0_.

To further summarize the results and to formally check for the presence of an inflection point, we fit to our simulated data a function that has been previously used to phenomenologically model Allee-effect success probabilities (Drake 2004): the cumulative distribution function of the Weibull distribution

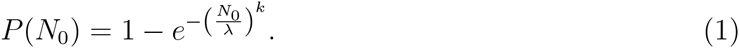

For different choices of the scale parameter *λ* and the shape parameter *k*, (1) can represent a wide range of success probability curves (Fig. S1). For *k* > 1, (1) represents the success probability under a strong Allee effect with critical population size

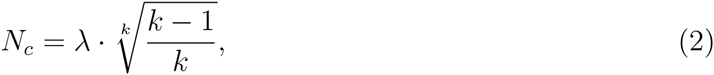

which is the founder population size at which the second derivative of *P*(*N*_0_) vanishes. For *k* ≤ 1, *P*(*N*_0_) does not have such an inflection point and therefore (1) can also be used to represent success probabilities in the absence of an Allee effect. In fact, for *k* = 1, (1) is of the form *P*(*N*_0_) = 1 − *u*^*N*_0_^ with constant *u*. This is the formula for the survival probability in a branching process, a model in which individuals produce offspring independently of each other and according to the same distribution (Karlin & Taylor 1975). Therefore, (1) is a natural null model for the detection of Allee effects.

To fit (1) to the success probabilities estimated from the simulations, we used the nonlinear least squares method as implemented in the “nls” function in R (R Core Team 2014). In some cases, the estimated value of *k* was above one, but the inflection point was at small population sizes and small success probabilities such that the success-probability curve was not markedly sigmoid. To highlight those cases that are biologically more relevant, we say that there is a “meaningful strong Allee effect” if the success probability curve has an inflection point and if the success probability at this inflection point is at least 1/3.

## Results

One after another, we will now consider the results for the four sets of simulations (**D, D+E, D+G, D+E+G**). We start with the case of only demographic stochasticity (**D**). Previously, Dennis (2002) showed that demographic stochasticity alone does not produce a strong Allee effect. This finding is generally confirmed by our simulation results (Fig. 1 a). Success probabilities rapidly increased with increasing founder population size. The success-probability curves were concave in the entire simulated range. The shape parameter, *k*, of the Weibull model was estimated to be slightly larger than one, but the resulting critical population sizes were below five, the smallest simulated founder population size, and also associated with a very low success probability, such that there was no meaningful Allee effect. A comparison with a hermaphroditic model version (Appendix S2) suggests that the small deviation from the expectation of *k* = 1 is due to random fluctuations in the sex ratio, which can cause extinction of very small populations in the two-sex model.

**Figure 1:**
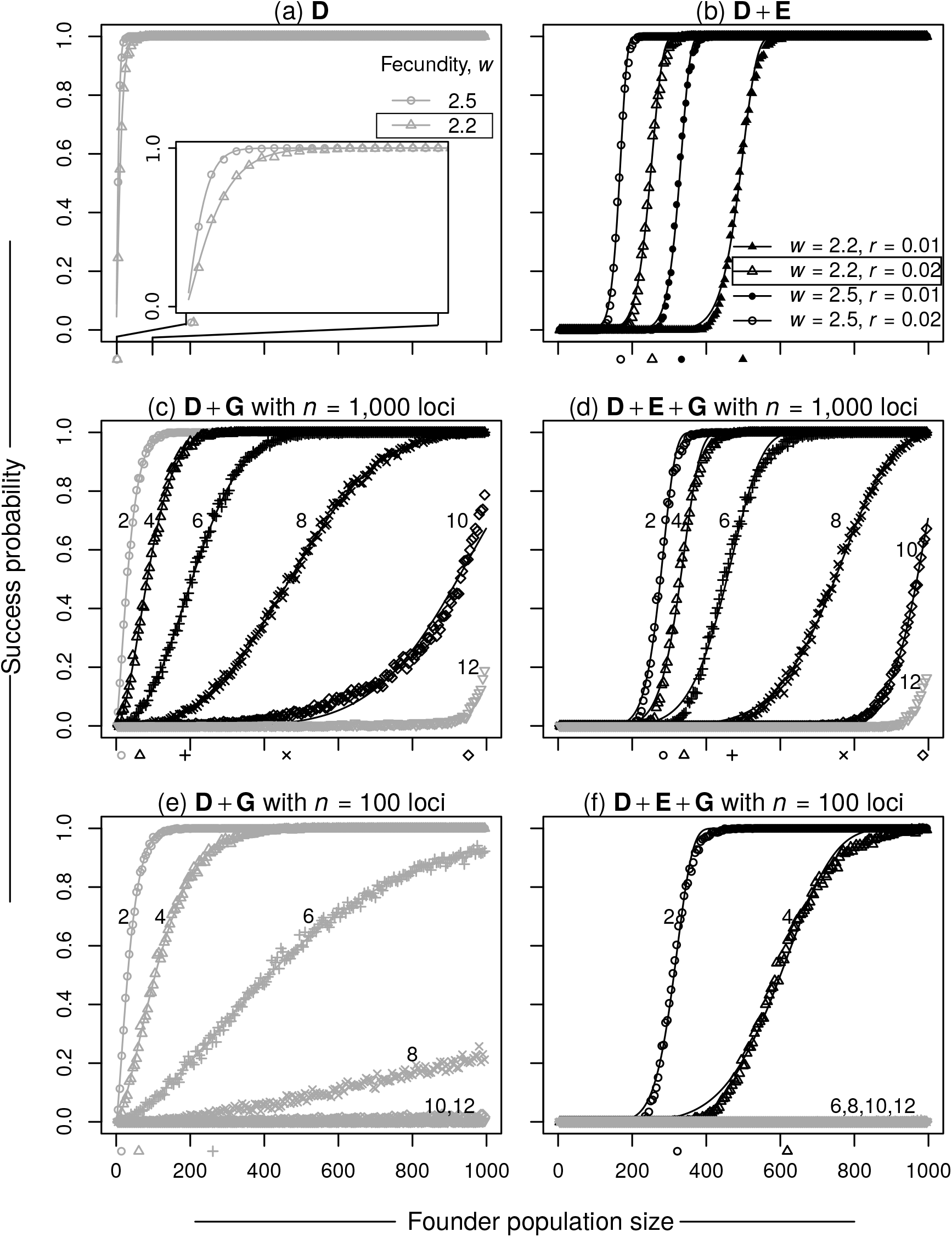
Example success-probability curves for the four sets of simulations. (a) Neither mate-finding problems nor genetic problems (the inset magnifies the results for founder population sizes up to 100). (b) Mate-finding problems, but no genetic problems. (c, e) Genetic problems, but no mate-finding problems. (d, f) Both mate-finding and genetic problems. The numbers next to the curves in (c–f) indicate the respective number of lethal equivalents, *b*, distributed either across *n* = 1, 000 loci (c, d) or across *n* = 100 loci (e, f). Symbols represent the proportion of successes in the simulations and lines are the corresponding fits of the Weibull model (1). Symbols below the *x*-axes indicate critical population sizes estimated from the Weibull model. Parameter combinations exhibiting a “meaningful” strong Allee effect (success probability at the critical size larger than 1/3) are shown in black, other parameter combinations in gray. For parameter combinations that had a success probability below 0.5, no model fit was attempted (also shown in gray). The boxed parameter combinations in (a) and (b) are those underlying (c–f).

Also as expected, mate-finding problems in the absence of genetic problems (**D+E**) did lead to clearly sigmoid success-probability curves and therefore meaningful strong Allee effects (Fig. 1 b). The position of the inflection point, i.e. the critical population size, increased with decreasing encounter rate, *r*, and with decreasing fecundity, *w*.

For populations with only genetic problems (**D+G**), the shape of the success-probability curve strongly depended on the number of lethal equivalents, *b*, i.e. the average number of lethal mutations carried by individuals in the source population, and on the number of loci that supplied these lethal mutations (Fig. 1 c and e, Fig. 2). With 1,000 loci, we obtained meaningful genetic Allee effects with clearly sigmoid success probability curves for sufficiently large values of *b*, approximately 4 to 6 depending on *w* (Fig. 1 c, Fig. 2 a). For smaller values of *b*, e.g. 2, the model fit indicated a small critical population size, but the curve was not strongly S-shaped and the Allee effect was not meaningful according to our definition since the success probability at the estimated critical size was very low. For very large numbers of lethal equivalents, success probabilities were low over the entire simulated range and no critical population size was estimated.

**Figure 2:**
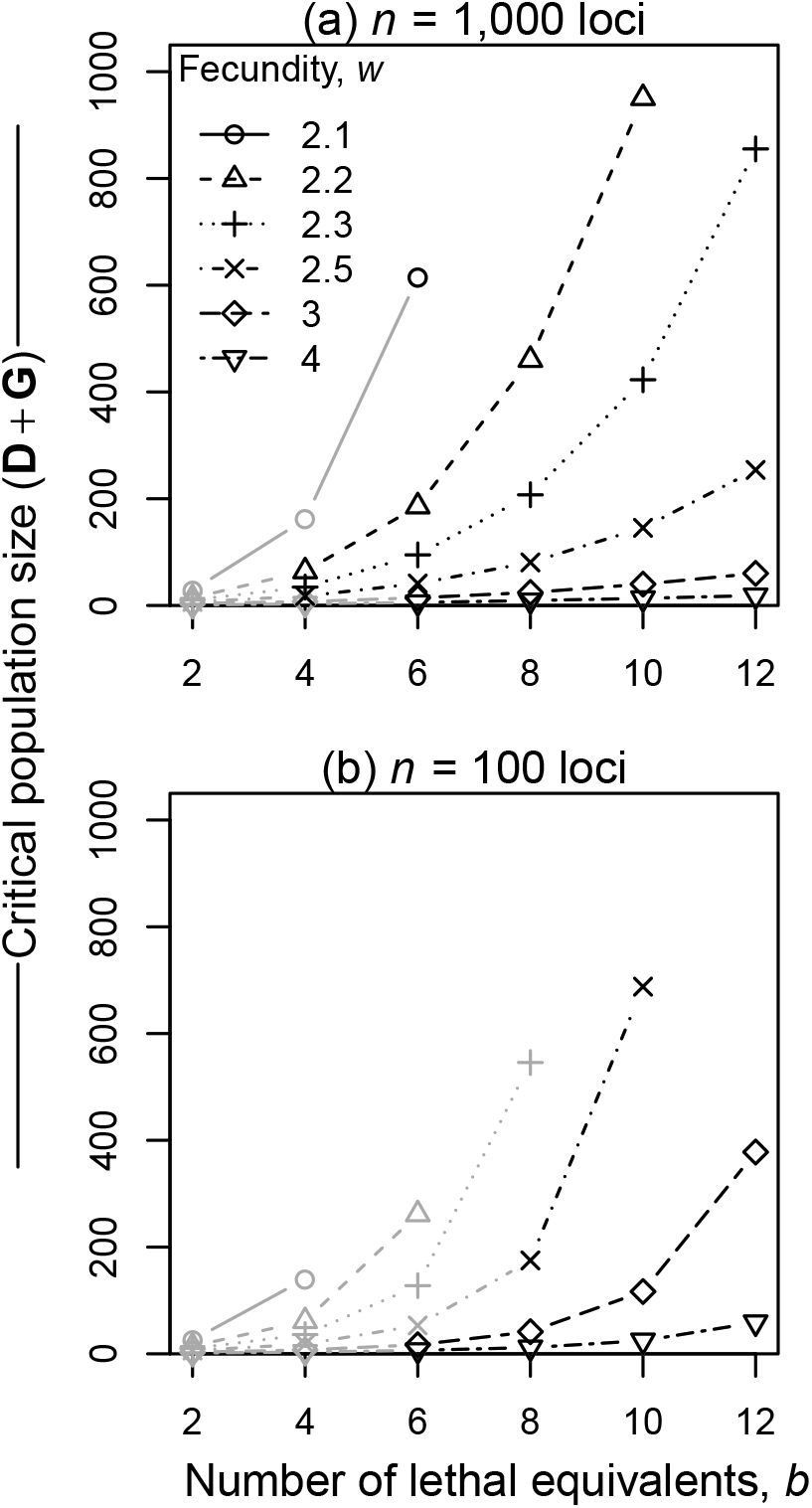
Critical population size with only genetic problems (**D+G**) with (a) *n* = 1, 000 loci or (b) *n* = 100 loci as a function of the number of lethal equivalents in the source population, *b*, for various values of the fecundity, *w*. Parameter combinations that exhibit a “meaningful” strong Allee effect (success probability at the critical size larger than 1/3) are shown in black, other parameter combinations in gray. Missing points correspond to cases where the Weibull model was not fit due to overall very low success probabilities.

For the same number of lethal equivalents, success probabilities were substantially lower with 100 contributing loci than with 1,000 loci (compare Fig. 1 e to Fig. 1 c). This reduction in success probability occurred over a broad range of founder population sizes, such that the S-shape was less pronounced than with 1,000 loci and our criterion for a meaningful strong Allee effect was often not fulfilled. However, in some cases with high numbers of lethal equivalents and high fecundity, meaningful strong Allee effects still emerged (Fig. 2 b). Both with 1,000 and with 100 contributing loci, estimated critical population sizes increased with the number of lethal equivalents, *b*, and with decreasing fecundity, *w* (Fig. 2). Also, despite large differences in overall success probabilities, the critical population sizes estimated for 100 and 1,000 loci were closely correlated (Fig. S2). Additional simulations with 5,000 loci yielded very similar results as for 1,000 loci, but critical population sizes were slightly lower (Fig. S3). As expected, critical sizes were insensitive to the choice of target population size (Fig. S4).

To elucidate the eco-evolutionary dynamics underlying the various success-probability curves for the **D+G** scenario, we considered the average population size trajectories and the average numbers of lethal alleles per individual in populations that are successful in the end (Fig. 3 a-f, see Fig. S5 for the corresponding results for failed populations). For few lethal equivalents, the average population size of successful populations increased monotonically over time (Fig. 3 a). For a larger number of lethal equivalents, non-monotonic population-size trajectories were common (Fig. 3 b, c), but took different forms depending on the underlying number of loci.

**Figure 3:**
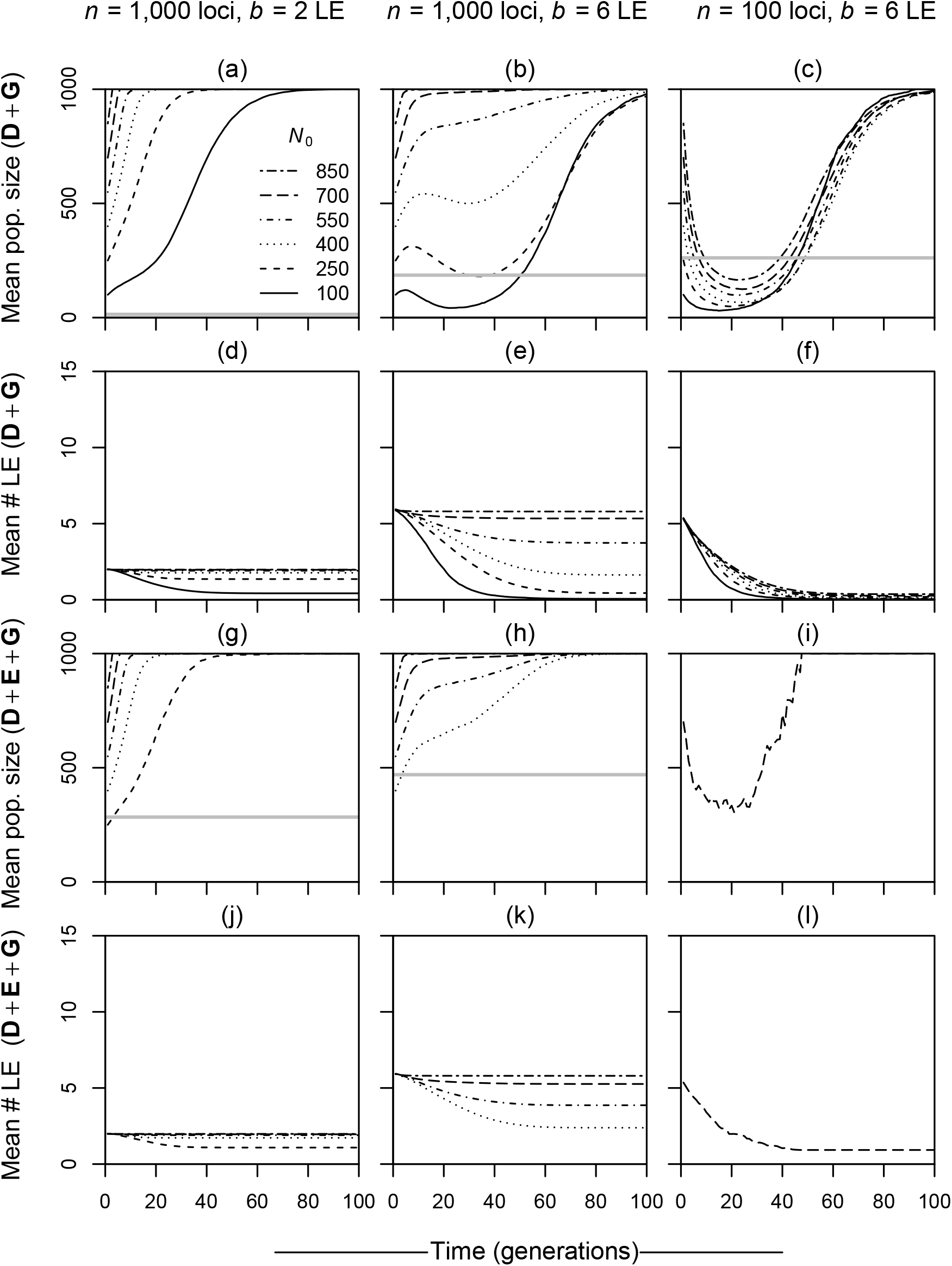
Eco-evolutionary dynamics for three example parameter combinations (one per column). (a–c) Average population size over time for successful **D+G** populations starting at different founder population sizes, *N*_0_. The horizontal gray lines indicate the critical population sizes estimated from the Weibull model. (d–f) Corresponding average numbers of lethal allele copies per individual over time for successful **D+G** populations. (g–l) Analogous results for successful **D+E+G** populations. Note that in some panels lines are missing because there were no successful populations for the respective founder population size. The corresponding results for failed populations are shown in Fig. S5. Other parameters: *w* = 2.2, *r* = 0.02.

With 1,000 loci, founder populations of small or intermediate size started to grow, then declined, often to numbers well below the founder population size, and eventually recovered (Fig. 3 b). Along the way, populations with small founder sizes purged almost all deleterious mutations. In fact, when reaching a population size of 1,000, they had on average fewer lethal mutations per individual than populations that had started with a smaller number of lethal equivalents (compare Fig. 3 e and d). By contrast, large populations often survived without substantially reducing the number of lethal mutations per individual (Fig. 3 e).

With 100 loci, lethal allele frequencies per locus were higher by design and therefore unrelated parents had a lower probability of producing viable offspring than with 1,000 loci. Consequently, in some cases all populations declined to low numbers, no matter how large they were initially (Fig. 3 c). Successful populations of all initial sizes purged their lethal alleles almost entirely before recovering (Fig. 3 f). For all values of *b* and *n*, mean allele frequencies decreased monotonically over time, while the variance across loci first increased and then decreased again (Fig. S6). The smaller the founder population size, the larger was the maximum variance in allele frequency.

Finally, we considered scenarios with ecological and genetic problems acting in the same population (**D+E+G**). For this, we combined the mate-finding Allee effect with an encounter rate, *r*, of 0.02 with various genetic Allee effects. Success probabilities were lower than in the corresponding other sets of simulations and critical population sizes were higher (Fig. 1 e, f). In many cases, however, success probabilities were so low over the entire range of founder population sizes, that no critical population size could be estimated.

With 1,000 loci, the critical population size with both ecological and genetic problems acting closely matched the sum of the two individual critical population sizes (Fig. 4 a). According to the terminology of Berec *et al*. (2007), ecological and genetic Allee effects are additive in these cases. This additivity was maintained when we increased the number of contributing loci to 5,000 (Fig. S3). In contrast to the case with just genetic problems (**D+G**, see Fig. 3 b), successful populations experiencing both genetic and ecological problems did not usually have non-monotonic population size trajectories (Fig. 3 h). It appears that the presence of an ecological Allee effects prevented the recovery of small populations via purging. Indeed, successful populations still harbored a substantial number of lethal mutations when reaching population size 1,000 (Fig. 3 k).

**Figure 4:**
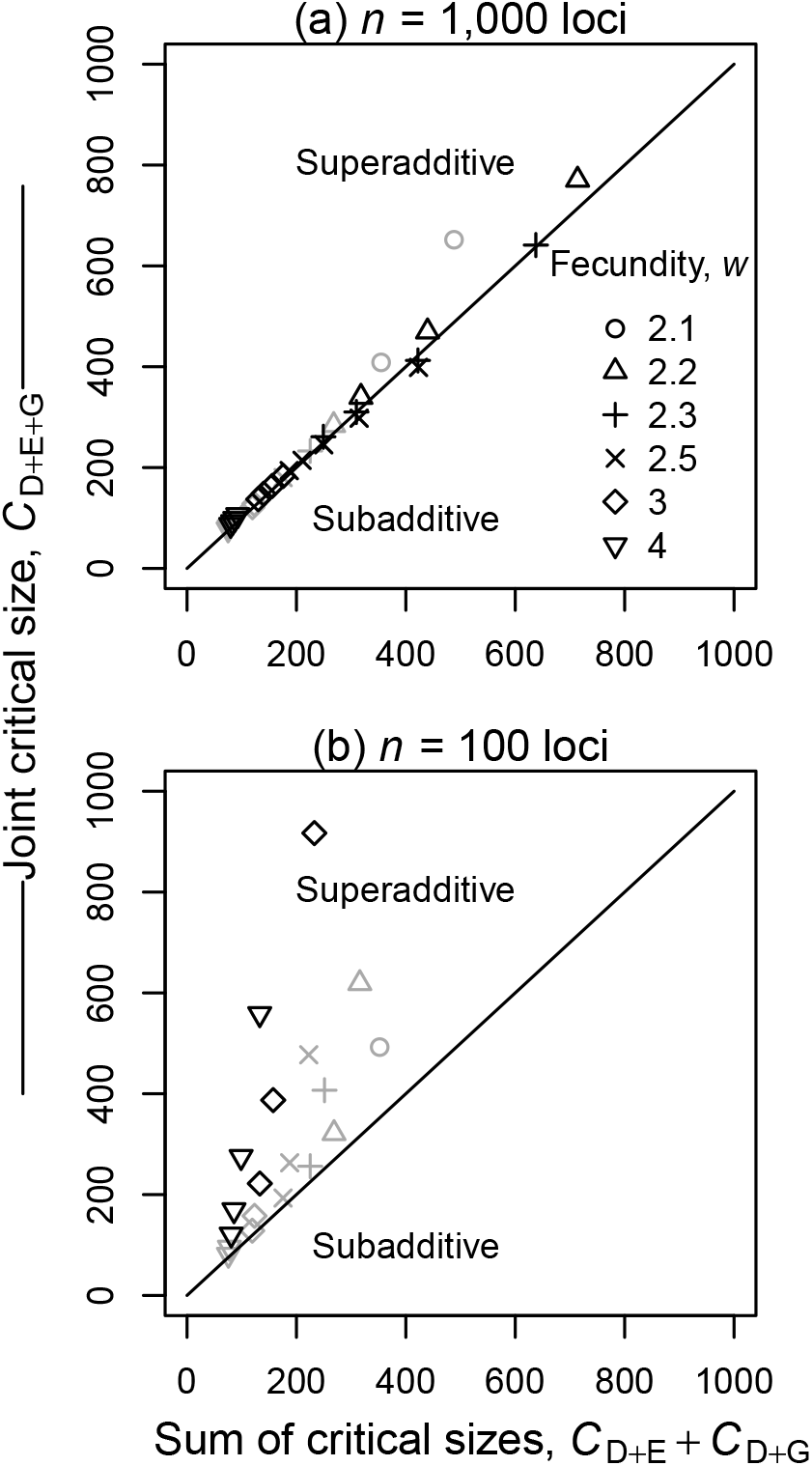
Relationship between the critical population size under both ecological and genetic problems, *C_D+E+G_*, and the sum of the critical sizes with only ecological problems acting, *C_D+E_*, or only genetic problems acting, *C_D+G_*. The lethal mutations are distributed either across *n* = 1,000 loci (a) or *n* = 100 loci (b). Critical population sizes are estimated by taking the inflection point of the fitted Weibull model (1). The various points for each number of loci correspond to different values of the fecundity, *w*, and the number of lethal equivalents, *b* (those represented in Fig. 2). The diagonal lines represent points where the Allee effects are additive. Parameter combinations that exhibit a “meaningful” strong Allee effect (success probability at the critical size larger than 1/3) are shown in black, other parameter combinations in gray. Fixed parameter: *r* = 0.02.

With 100 loci, ecological and genetic problems interacted in a qualitatively different manner. The critical population size under both effects was generally larger than the sum of the individual critical population sizes (Fig. 4 b). Thus the Allee effects were superadditive according to the terminology of Berec *et al*. (2007). Even when genetic problems alone did not lead to a meaningful strong Allee effect, they could strongly increase the critical population size when combined with mate-finding problems. A good example is the case of four lethal equivalents in Fig. 1. In this case, the success probability at a founder population size of around 400 was close to one with either only ecological problems (Fig. 1 b) or only genetic problems (Fig. 1 e) acting, but close to zero with both acting at the same time (Fig. 1 f).

To explore the robustness of our results and their dependence on mating system and genetic architecture, we have studied a number of alternative model versions. The results were generally very robust to changes in the mating system. Both a model version where females could mate with multiple males and a hermaphroditic model version yielded qualitatively similar results for individual Allee effects as well as their interaction (Appendix S1 and S2, Figs S8 and S9). The only major difference was that critical population sizes under genetic problems were slightly lower than in the original model. This result is expected because both multiple mating events and hermaphroditic reproduction lead to a higher number of distinct couples that contribute to the next generation and therefore relatedness and inbreeding depression should be reduced.

Further simulations of an extended model version (Appendix S3) indicate that the results are more sensitive to assumptions about the genetic architecture. Given the same average number of lethal alleles in the founder population, changing the dominance coefficient from 0 to 0.05 overall reduced success probabilities, but also led to smaller critical population sizes and in many cases extinguished the signature of a strong Allee effect (Fig. S10). When we assembled a given number of lethal equivalents from the empirical distribution of selection and dominance effects estimated for yeast (Agrawal & Whitlock 2011), meaningful genetic Allee effects emerged for appropriate choices of the fecundity, *w* (Fig. S12). As in the original model, successful populations often had non-monotonic population size trajectories. The interaction with an ecological Allee effect was superadditive (Fig. S12 c). When we assumed that half of the lethal equivalents are due to fully recessive lethals and the other half due to fully recessive mutations of smaller effect (Fig. S13), clear genetic Allee effects with sharp extinction thresholds emerged. However, successful populations usually maintained a positive growth rate at all times. Populations that declined due to inbreeding depression usually failed to purge enough mutations and went extinct. Unlike in all other scenarios, the interaction of these genetic Allee effects with ecological Allee effects was subadditive (Fig. S13 b).

## Discussion

### Genetic Allee effects

Our results indicate that recessive lethal mutations can indeed lead to a meaningful strong Allee effect with a sharp increase in establishment probability around a critical population size, but only under certain conditions. Most importantly, the average number of lethal mutations per haploid genome in the source population (lethal equivalents) needed to be larger than around five. Given that the average bird or mammal population has been estimated to carry approximately six haploid lethal equivalents (O’Grady *et al*. 2006), we expect that a large proportion of species will be susceptible to such strong genetic Allee effects.

For the same number of lethal equivalents, we found qualitatively different eco-evolutionary dynamics depending on whether the lethal equivalents were distributed across 1,000 or 100 loci. With 1,000 loci, lethal allele frequencies at each locus were small enough initially so that small founder populations of initially unrelated individuals started to grow. Eventually relatedness in small populations increased and inbreeding depression took effect (see Fig. 3). While populations with small founder size declined to extinction, larger populations usually managed to purge their deleterious mutations and eventually recovered. With 100 loci, by contrast, lethal allele frequencies per locus were relatively high to start with. Consequently, even unrelated founder individuals failed to produce enough viable offspring and populations of any size declined from the start. If founders could be assumed to stem from a large source population at equilibrium and under the same selection pressures as the newly founded population, high lethal allele frequencies would be implausible. However, high initial lethal frequencies might occur if the founding event or reduction in population size is accompanied by a sudden change in the environment such that mutations that were previously neutral or slightly deleterious are now lethal. Such an emergence of new lethal or almost lethal mutations has been demonstrated for example in *Drosophila melanogaster* under high temperature and other stressful conditions (Bijlsma *et al*. 1999). Populations would then need to adapt to the altered conditions, i.e. purge the newly lethal mutations, to escape extinction. This is a form of evolutionary rescue (Gomulkiewicz & Holt 1995; Gonzalez *et al*. 2013). Therefore, we may call the genetic Allee effect at 100 loci an “evolutionary-rescue genetic Allee” as opposed to the “inbreeding-depression genetic Allee effect” observed at 1,000 loci.

More generally, our results suggest that with an increasing number of loci contributing to inbreeding depression, success probability curves become more and more sigmoid with increasingly sharp extinction thresholds (compare e.g. Fig 1 c and e, Fig. S13). Lethal equivalents concentrated at relatively few loci can strongly increase extinction risk, but they do so over a wide range of founder population sizes. Consequently, there is no clear extinction threshold. In Appendix S4, we consider the extreme case of fully recessive lethal mutations at a single locus and derive an analytical approximation for the success probability function. As for the model with just demographic stochasticity, the success probability can be written in the Weibull function form of (1) with *k* =1. This result indicates that fully recessive lethal mutations at a single locus, no matter how frequent they are, never cause a strong genetic Allee effect. The key intuition underlying this result is that, since heterozygotes have the same fitness as wildtype homozygotes, reproduction of wildtype lineages is in no way affected by the presence of lethal mutant alleles in the population.

Both the inbreeding-depression genetic Allee effect and the evolutionary-rescue Allee effect were characterized by non-monotonic population-size trajectories for a range of founder population sizes (see Fig. 3). This makes the two genetic Allee effects qualitatively different from simple ecological Allee effects where growth rate just depends on population size and non-monotonicity is consequently not possible (it would imply two different growth rates at the same population size). We conjecture, however, that more complex ecological scenarios involving spatial structure or feedbacks with other species could in principle also cause an Allee effect with non-monotonic population-size trajectories. For example, classical predator-prey theory (see e.g. Begon *et al*. 2006) predicts that prey populations of the same size may have different per-capita growth rates depending on the population size of the predator. If a prey population is suddenly reduced in size, it will generally take some time for the predator population size to respond. In the meantime, the prey population will suffer from an increased predation risk and further decline in size. Eventually, the predator population size will also shrink and the prey population may be able to recover.

### Interaction between ecological and genetic Allee effects

When multiple strong Allee effects due to different mechanisms act in the same population, one might expect that a population will survive if it is above the largest of the critical population sizes of the individual Allee effects. This would imply a subadditive interaction between Allee effects, where the joint critical population size is smaller than the sum of the individual critical sizes. Contrary to this expectation, our results indicate that the interaction of a genetic Allee effect with a mate-finding ecological Allee effect is generally either additive or superadditive (see Fig. 4). That is, populations with founder sizes well above the critical sizes of both ecological and genetic Allee effect still went extinct when both acted together. This phenomenon can be explained by the non-monotonicity of population-size trajectories under a genetic Allee effect (see Fig. 3). Over a wide range of founder population sizes, populations declined to sizes well below the founder size. In the absence of an ecological Allee effect, these reduced populations could purge their deleterious mutations and recover. However, in the presence of an ecological Allee effect such populations were driven to extinction by mate-limitation before the deleterious mutations could be purged. Thus the ecological Allee effects prevented populations from purging their deleterious mutations. Indeed, successful populations under both Allee effects harbored more deleterious mutations than successful populations under just the genetic Allee effect.

With 1,000 loci (and also with 5,000 loci), interactions between ecological and genetic Allee effects were strikingly close to additive (see Fig. 4 a, Fig. S3 b). With 100 loci, the interaction was generally superadditive (see Fig. 4 b). Even if the critical population size was small with only genetic problems acting, genetic problems caused large increases in critical population size when added to a population already suffering from mate-finding difficulties. In Fig. 5, we offer a heuristic explanation for the emergence of additivity at 1,000 loci and superadditivity at 100 loci.

**Figure 5:**
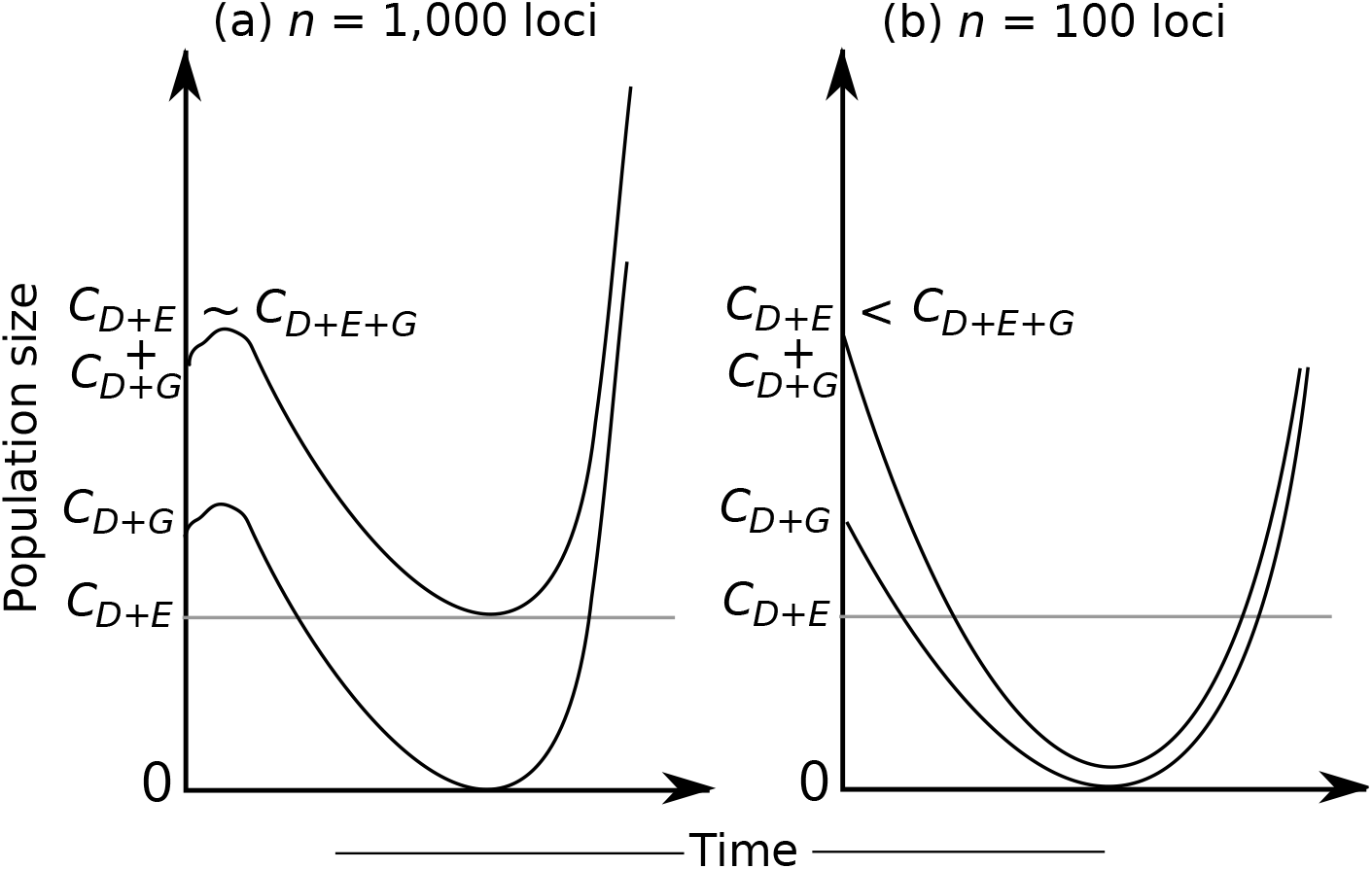
Intuitive explanation for (a) additivity of Allee effects with *n* = 1,000 loci and (b) superadditivity with *n* = 100 loci. *C_D+E_, C_D+G_*, and *C_D+E+G_* respectively are the critical sizes with only ecological problems, only genetic problems, or both. In each panel, the curves represent two typical population-size trajectories for populations with only a genetic Allee effect (**D+G**), one starting at founder population size *C_D+G_* and the other at *C_D+G_* + *C_D+E_*. Since *C_D+G_* is the critical population size, a typical trajectory starting at this founder population size almost reaches population size zero at its lowest point, if it is successful at all. In the case of 1,000 loci (a), the simulation results (see Fig. 3 b) suggest that over a range of founder population sizes, typical trajectories for different founder sizes are merely shifted by a constant. Therefore, populations starting at size *C_D+G_* + *C_D+E_* will have approximately size *C_D+E_* as the lowest point. Now we add an ecological Allee effect and assume that it only affects populations once they are close to its critical size, *C_D+E_*. We see that *C_D+G_* + *C_D+E_* is the founder population size for which a population barely escapes the ecological Allee effect and therefore extinction. Thus, *C_D+E+G_* ≈ *C_D+G_* + *C_D+E_*. In the case of 100 loci (b), population-size trajectories for different founder population sizes did not merely differ by a constant. Over a wide range of founder population size, populations first crashed to very small sizes because of large lethal allele frequencies (see Fig. 3 c). Therefore, populations with founder size *C_D+G_* + *C_D+E_* typically go extinct if the ecological Allee effect is added. Thus *C_D+E+G_* > *C_D+G_* + *C_D+E_*.

If we were to examine values for the number of loci between 1,000 and 100, we would expect increasingly superadditive interactions with decreasing number of loci. In fact, even with 1,000 loci the interaction appears to be not exactly additive, but very slightly superadditive (see Fig. 4 a). For numbers of loci beyond 1,000 or 5,000, we would expect the interaction to remain additive. Already with 1,000 loci, it is quite unlikely that multiple individuals in the founder population carry a mutation at the same locus. If we increase the number of loci while keeping the number of lethal equivalents constant, we eventually reach a point where every lethal mutation is unique in the founder population, at which point further increases in the number of loci have no effect.

Given that two Allee effects in our study generally interacted additively or superadditively, one might wonder whether this pattern will hold up with more than two Allee effects acting in the same population. For example, there could be an additional predator-induced Allee effect or an Allee effect due to a different genetic problem, such as mutational meltdown. We conjecture that further Allee effects associated with non-monotonic population size trajectories will substantially increase the critical population size. By contrast interactions involving multiple Allee effects with monotonic population size trajectories will supposedly be subadditive. This is supported by our observation that ecological and genetic Allee effects were subadditive when half the lethal equivalents were due to recessive mutations of small effects that apparently could not be purged, such that population-size trajectories were generally monotonic. Exploring these hypotheses is an interesting direction for future work, especially given that many small populations face multiple challenges, both ecological and genetic.

### Robustness and potential limitations

In this study, we have assumed for simplicity that all loci recombine freely, i.e. that there is no linkage between loci. This may appear unrealistic if the number of loci is on the order of thousands and thus the average genetic distance between loci must be relatively small. However, both in our simulations and in the empirical studies that motivated our parameter choices (see Table 1 in O’Grady *et al*. 2006) the average number of lethal mutations carried by an individual was at most on the order of tens. Therefore, loci with mutations tend to be far apart in the genome and segregate almost independently. In any case, closely linked lethal mutations essentially behave like lethal mutations at a single locus. Hence, also the empirically-derived estimates for the number of lethal equivalents represent an effective number of unlinked mutations rather than the actual number of mutations. Thus, the assumption of free recombination appears generally justified.

We have further assumed that generations are discrete. Overlapping generations may substantially increase the survival prospects of small populations because long-living adults can attempt multiple times to find a partner and produce viable offspring. In some species of invasive plants, small populations can even reproduce clonally until sufficient pollen of the right genetic types becomes available (McCormick *et al*. 2010). However, it is currently not clear whether such effects only increase overall success probability or whether they change the qualitative shape of the success-probability curve.

Our explorations of alternative model versions indicate that the presence of genetic Allee effects is robust to changes in the details of the mating system (see Appendix S1 and S2), but is more sensitive to the genetic architecture underlying a given number of lethal equivalents (see Appendix S3). Nevertheless, strong genetic Allee effects emerged for suitable parameter combinations under all genetic architectures we considered. In particular, the most complicated and most realistic architectures produced some of the clearest cases for strong genetic Allee effects (see Figs S12 and S13).

### Implications for endangered and introduced populations

Although our models are too simplistic to derive specific recommendations for the management of endangered or introduced populations, we can highlight a few areas where an understanding of genetic Allee effects and their interactions with ecological Allee effects may be helpful. First, neglecting the possibility of a genetic Allee effect may cause us to make overly optimistic predictions for the fate of an endangered species. For instance, if a small population grows after a population crash, we might assume that it will keep growing and quickly recover to its previous size. However, our results suggest that populations with a strong genetic Allee effect often grow initially but then decline to extinction due to inbreeding depression.

Our results further suggest that it can be very important to account for the interaction of ecological and genetic problems when assessing the viability of endangered populations. As an example, imagine we are worried about the fate of a particular population that has just been decimated to approximately 450 individuals by a severe drought. We now independently carry out two analyses. First an analysis of mating success and other ecological variables tells us that a population size of approximately 200 individuals is necessary to avoid extinction from ecological problems. Second a genetic analysis informs us that about 300 individuals would be enough to avoid extinction due to inbreeding depression and other genetic problems. Since the population is well above either of these critical sizes we might come to an optimistic prognosis. However, our simulation results indicate that critical population sizes of ecological and genetic Allee effects are generally additive or even superadditive, and thus that the combined critical population size will be around or above 500. Therefore, our population may in reality be at high risk of extinction and active protection measures should be taken. One possibility would be to introduce individuals from carefully selected populations at other locations or even from other subspecies (Hedrick & Kalinowski 2000), as has been done for example for the Florida panther (Johnson *et al*. 2010).

Finally, taking into account genetic Allee effects and their interaction with ecological Allee effects may help with intentional introductions of biological control agents or with re-introductions of extinct populations. For example, previous work has shown that in the presence of a strong Allee effect and given a fixed number of individuals available for introduction, it is better to make few large introductions rather than many small introductions (Grevstad 1999; Shea & Possingham 2000; Wittmann *et al*. 2014c). In the presence of a genetic Allee effect, one should additionally try to select founder individuals from diverse genetic backgrounds.

### Future directions for research on genetic Allee effects

One important goal of our study was to contribute toward more rigorous criteria for genetic Allee effects and to clarify our thinking about them. Our definition via the inflection point of the success-probability curve provides a criterion for a strong genetic Allee effect that can be applied even if the relationship between population size and per-capita growth rate is confounded by changes in genetic composition. Also motivated by the need for clearer definitions of genetic Allee effects, Luque *et al*. (in press) recently proposed to define a genetic Allee effect as a two-step process, where a reduction in population size first leads to a change in the genetic structure of the population and, second, this change in genetic structure reduces a fitness component. An advantage of this definition is that it includes weak Allee effects and component Allee effects. However, it may sometimes be challenging to decide when exactly to assess genetic structure and fitness. As we have seen, the growth rate of a small founder population can be first positive, then negative, and finally positive again. Because of purging, the final growth rate can even be higher than in populations that were initially larger.

The study by Luque *et al*. (in press) and our study are only first steps in the exploration of genetic Allee effects. There are several important directions for future research, both empirical and theoretical. On the empirical side, an exciting but ambitious future direction is to document genetic Allee effects in the sense used in this study, i.e. demonstrate the existence of inflection points in the success-probability curve. Since such analyses require many replicate populations with a range of initial population sizes, they are virtually impossible for natural populations or even endangered species. But they might be feasible in laboratory experiments on organisms with short generation times such as *Drosophila*. For example, Kramer & Drake (2010) attempted such an analysis to study predator-mediated Allee effects in a *Daphnia-Chaoborus* system. However, it is important to carefully consider whether and how results on model organisms in the laboratory apply to endangered or introduced populations in the wild.

Another important goal is to improve our understanding of the genetic architecture of inbreeding depression in species of concern. Our simulation results indicate that even for the same number of lethal equivalents, a population’s survival prospects can depend strongly on the number of loci with deleterious mutations and the distribution of selection and dominance coefficients. Yet, these parameters are generally unknown. Often, we only know the average number of lethal equivalents for some other species in the same broad taxonomic group, e.g. birds or mammals (Ralls *et al*. 1988; O’Grady *et al*. 2006). This quantity is then used, for example as input for population viability analyses, combined with ad-hoc assumptions about the genetic architecture. Fortunately, detailed data on the distribution of selection and dominance effects are now becoming available for some model organisms, e.g. yeast (Agrawal & Whitlock 2011). Such studies are extremely labor intensive and require the systematic generation of mutants, such that they are certainly not feasible for actual species of concern. However, it would be very helpful to have results for a number of model organisms from different taxonomic groups. If the distributions of selection and dominance coefficients, for example, are consistent across groups, then we may be more confident in using them also to model the dynamics of endangered species.

On the theoretical side, important questions are whether and under what conditions loss of genetic variation and mutation accumulation, the two other main genetic problems of small population size, can also give rise to a strong Allee effect. As a next step, we are planning to explore the relationship between population size and the extinction risk of a population in a temporally fluctuating environment. Since ecological Allee effects can influence the amount of genetic variation maintained by a founder population (Kramer & Sarnelle 2008; Wittmann *et al*. 2014a,b), we expect to find interesting interactions between ecological Allee effects and these genetic problems.

## Acknowledgments

For discussion and comments, we thank members of the Biomathematics group at the University of Vienna, as well as Sebastian Matuszewski and Claus Vogl. MJW acknowledges fellowships from the Stanford Center for Computational Evolutionary and Human Genomics (CEHG) and from the Austrian Science Fund (FWF, M 1839-B29). Simulations were performed on the Vienna Scientific Cluster (VSC).

## Supplementary Information

**Figure S1:**
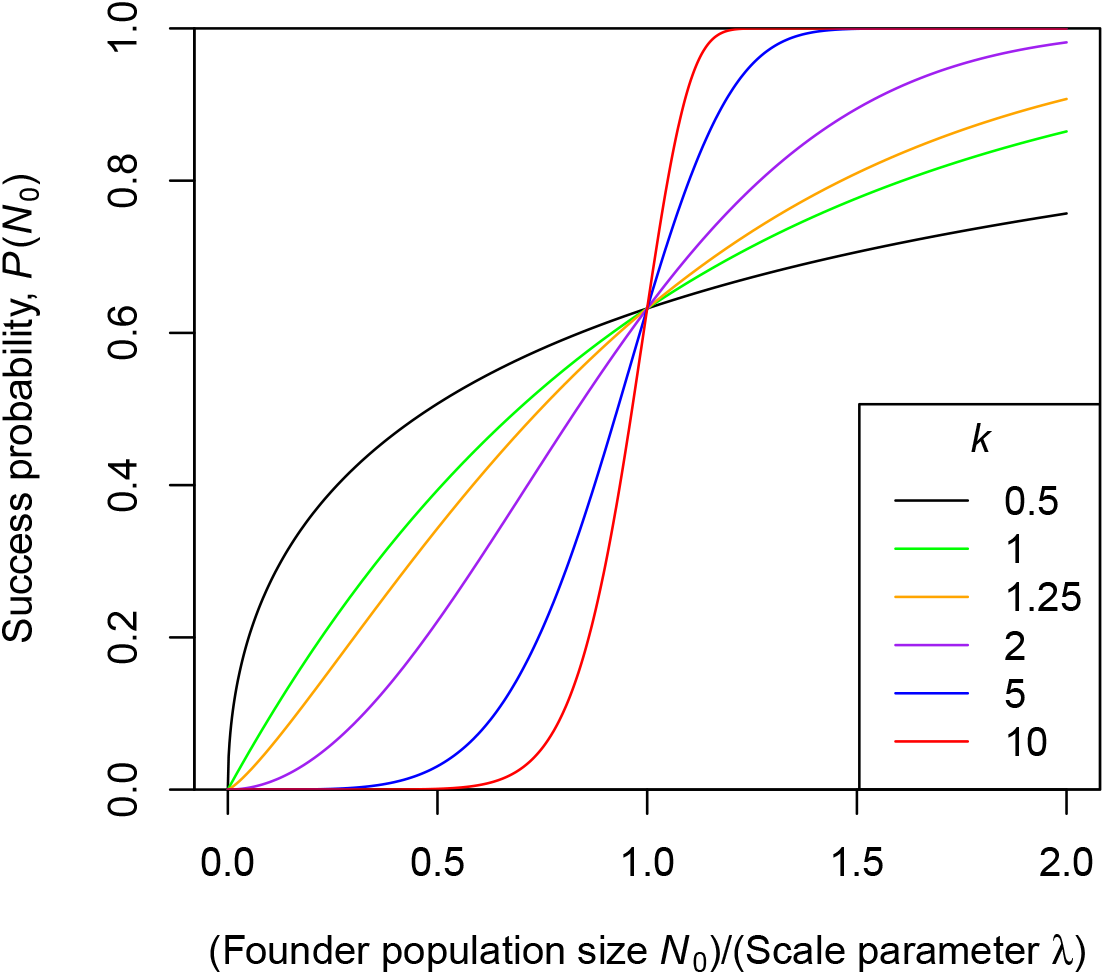
Examples for various shapes of the success probability curve that can be represented by the Weibull model (1).

**Figure S2:**
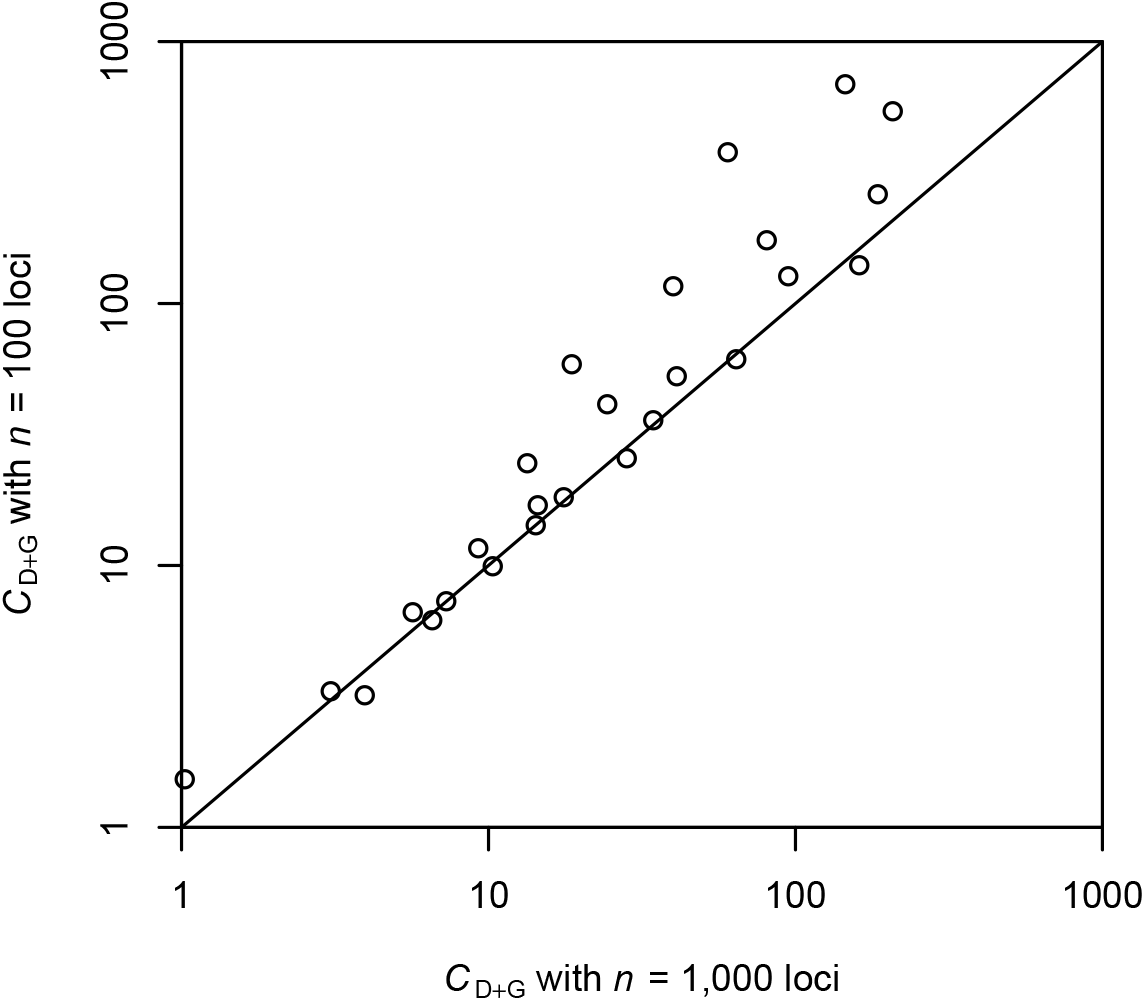
Critical population sizes with only genetic problems (**D+G**) at 100 loci vs. 1,000 loci. On the diagonal line, both critical sizes are equal.

**Figure S3:**
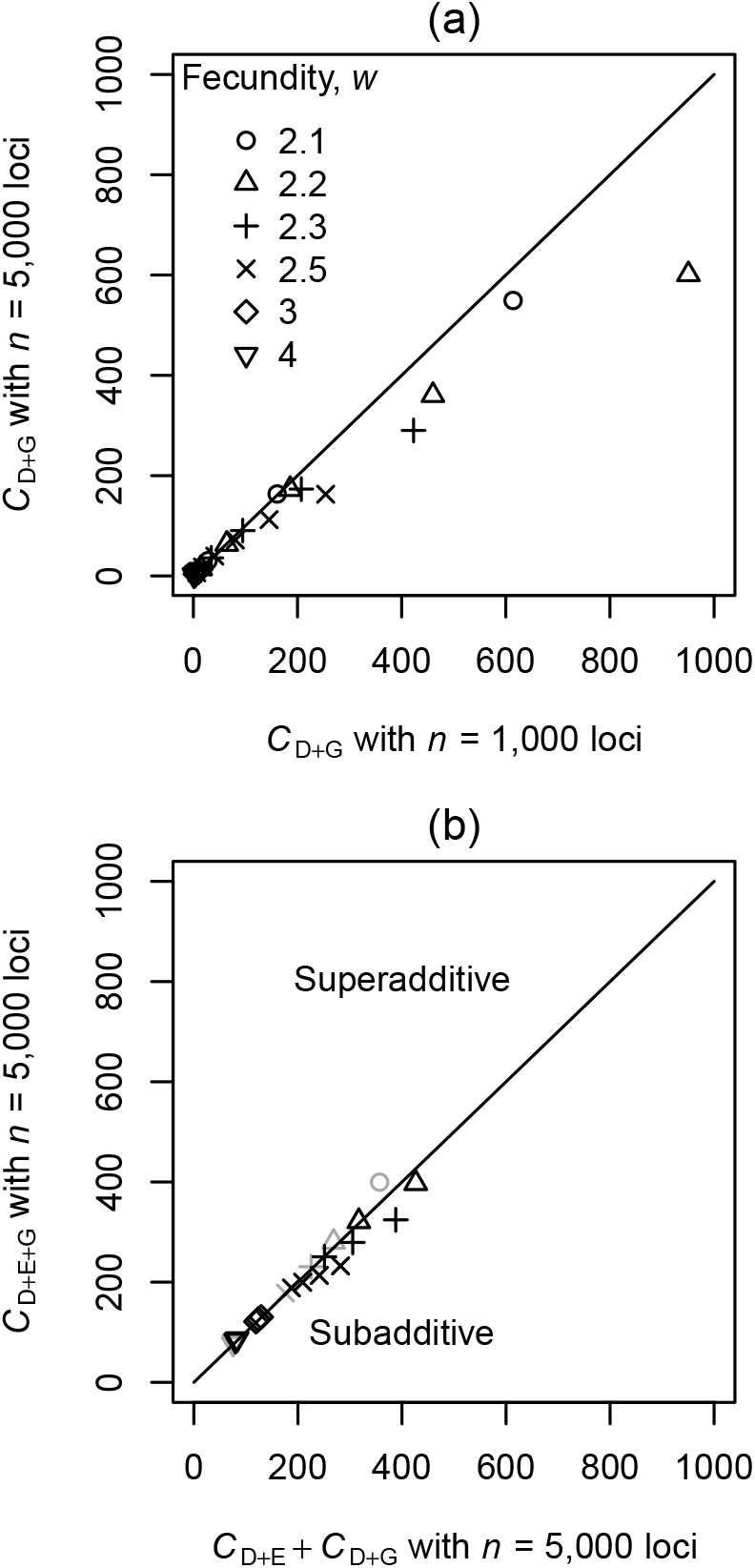
Summary of results with 5,000 loci contributing to inbreeding depression. (a) Comparison of critical population sizes with only genetic problems (**D+G**) and the same number of lethal equivalents spread over 5,000 loci vs. 1,000 loci. (b) Relationship between the critical population size under both ecological and genetic problems, *C_D+E+G_*, and the sum of the critical sizes with only ecological problems acting, *C_D+E_*, or only genetic problems acting, *C_D+G_*. Parameter combinations that exhibit a “meaningful” strong Allee effect (success probability at the critical size larger than 1/3) are shown in black, other parameter combinations in gray. In both (a) and (b), the various points correspond to different values of the fecundity, *w*, and the number of lethal equivalents, *b*. The diagonal lines represent points where both critical population sizes are equal (a) or the Allee effects are additive (b).

**Figure S4:**
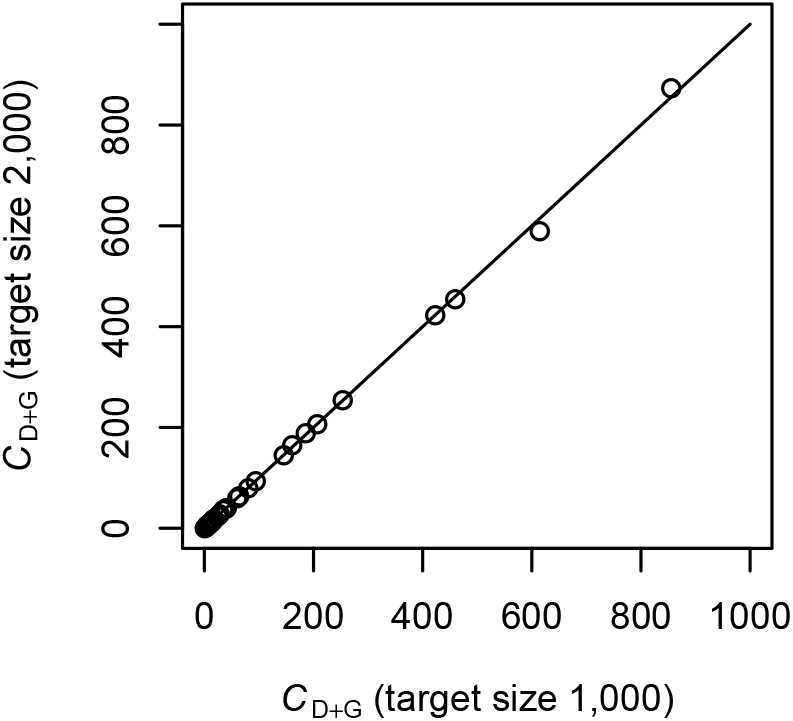
Critical population sizes with only genetic problems (**D+G**) with a target size of 2,000 compared to the default target size of 1,000. On the diagonal line, both critical sizes are equal.

**Figure S5:**
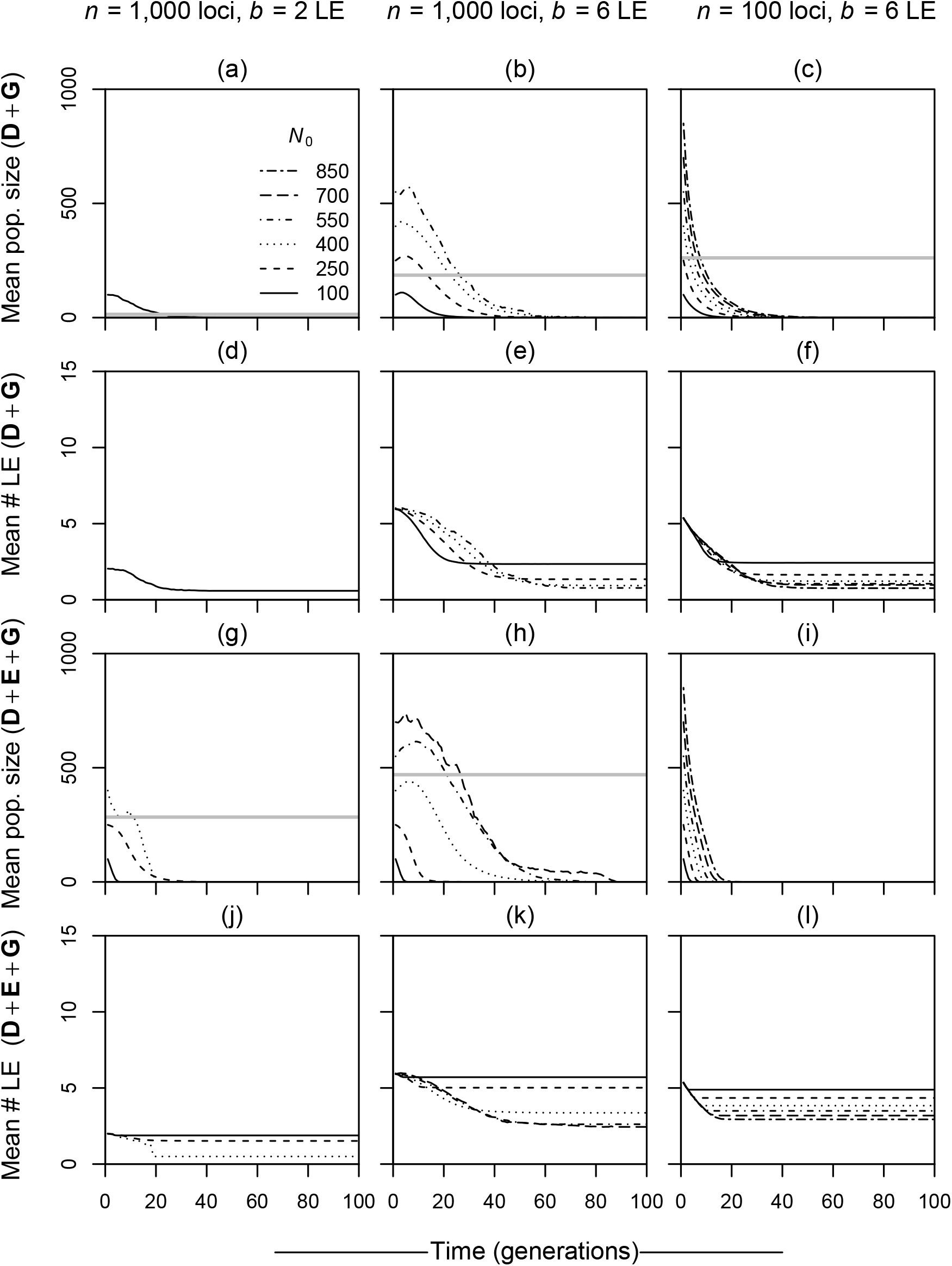
Eco-evolutionary dynamics of failed populations for three example parameter combinations (one per column). (a–c) Average population size over time for failed **D+G** populations starting at different founder population sizes, *N*_0_. The horizontal gray lines indicate the critical population sizes estimated from the Weibull model. (d–f) Corresponding average numbers of lethal allele copies per individual over time for failed **D+G** populations. (g–l) Analogous results for failed **D+E+G** populations. Note that in some panels lines are missing because there were no failed populations for the respective founder population size. The corresponding results for successful populations are shown in Fig. 3. Other parameters: *w* = 2.2, *r* = 0.02.

**Figure S6:**
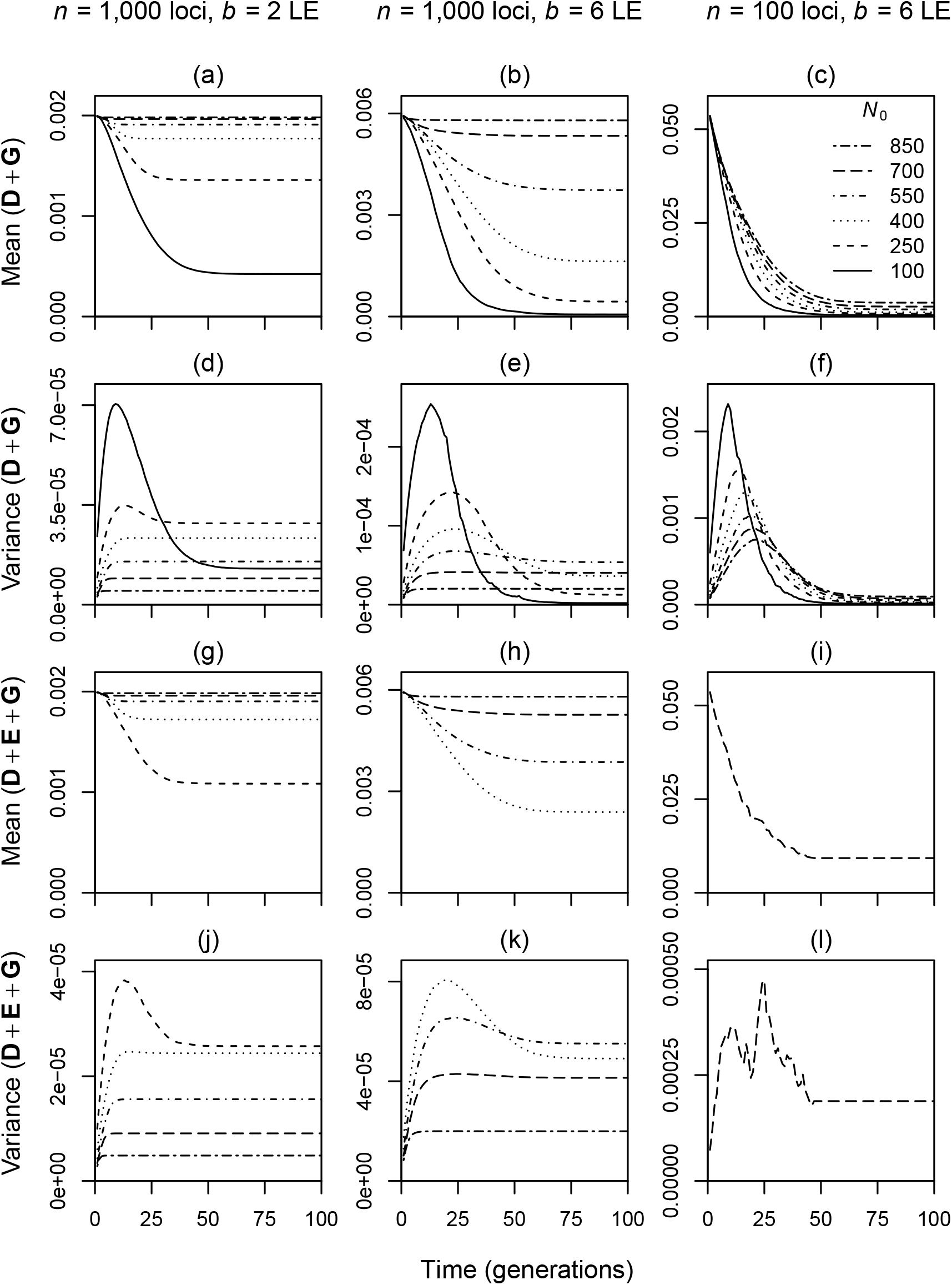
Dynamics of the mean allele frequency over loci (a–c, g–i) and the corresponding variance in allele frequency (d–f, j–l) for successful populations starting at different founder population sizes, *N*_0_ (see legend in c). (a–f) are for the **D+G** scenario and (g–l) for the **D+E+G** scenario. Note the different scales of the *y*-axes. The corresponding results for failed populations are shown in Fig. S7. Other parameters: *w* = 2.2, *r* = 0.02.

**Figure S7:**
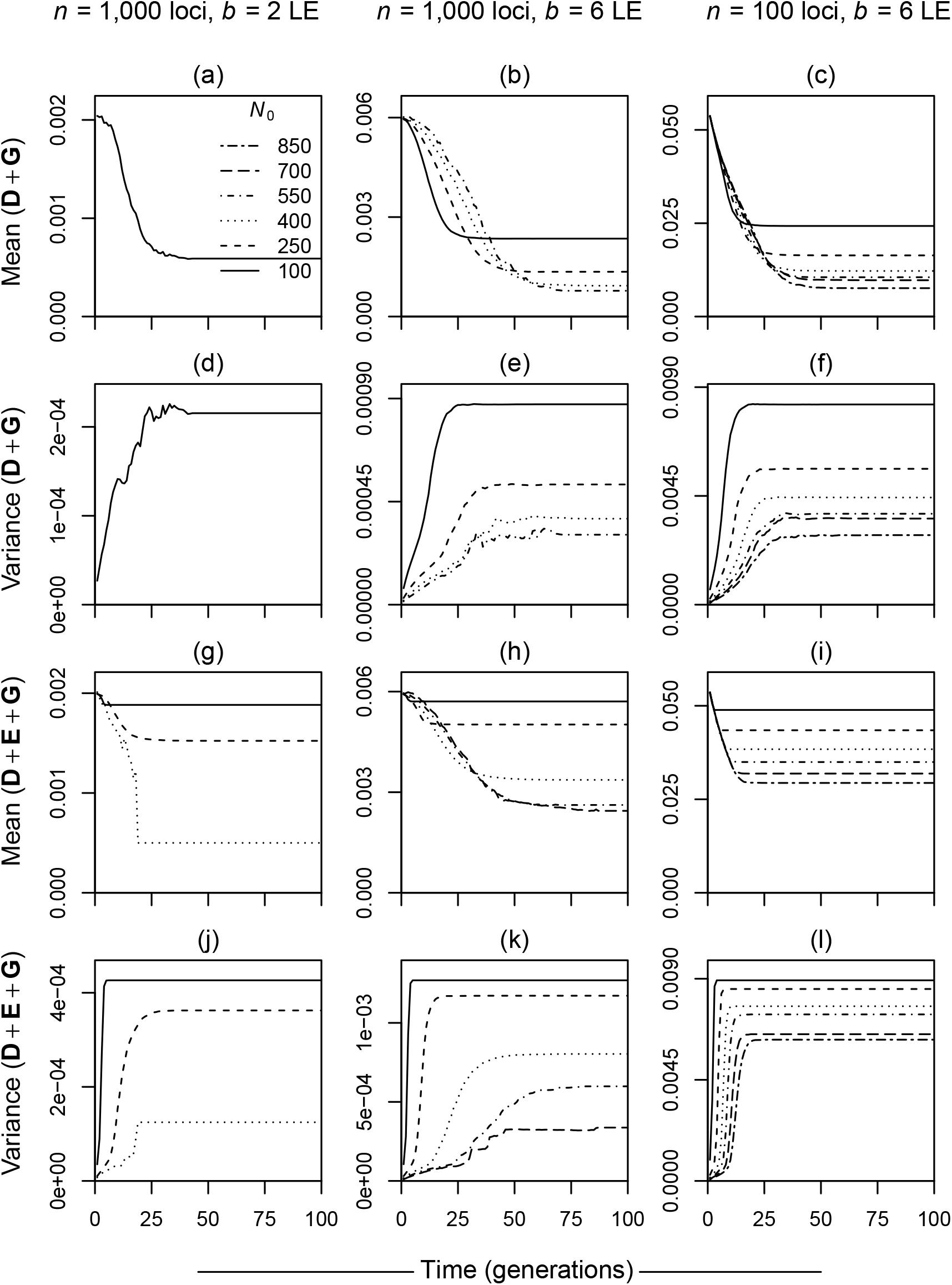
Dynamics of the mean allele frequency over loci (a–c, g–h) and the corresponding variance in allele frequency (d–f, j–l) for failed populations starting at different founder population sizes, *N*_0_ (see legend in a). (a–f) are for the **D+G** scenario and (g–l) for the **D+E+G** scenario. Note the different scales of the y-axes. The corresponding results for successful populations are shown in Fig. S7. Other parameters: *w* = 2.2, *r* = 0.02.

## Appendix S1 Model version with multiple mating events per female

Independently of all other females, each female in generation *t* encounters a Poisson-distributed number of potential mating partners with mean *r* · *M_t_*, where *r* is the per-capita encounter rate and *M_t_* is the number of males in generation t. We draw each encountered male independently and with replacement from all males in the population, which means that the same male can be encountered multiple times and then has a higher probability to sire offspring.

If a female finds at least one mating partner (probability 1 − *e*^−*rηM_t_*^), she produces a Poisson-distributed number of offspring, *X*, with mean *w*_0_, as in the original model. We thereby assume that the number of offspring produced by a female is independent of the number of mating partners, as long as she mated at least once. Each of the *X* offspring is independently assigned a father by randomly drawing one of the males that the mother mated with. In the simulation sets without mate-finding problems (**D** or **D+G**), each female effectively mates with all males, such that each offspring draws a father independently from all males in the population. In all other respects, the simulation proceeded as for the original model.

Compared to the original model with a single mating event per female, the model version with multiple mating events produced slightly lower critical population sizes under a genetic Allee effect (Fig. S8 a). But the results were very similar in all other respects, including the additivity of ecological and genetic Allee effects (Fig. S8 b).

**Figure S8:**
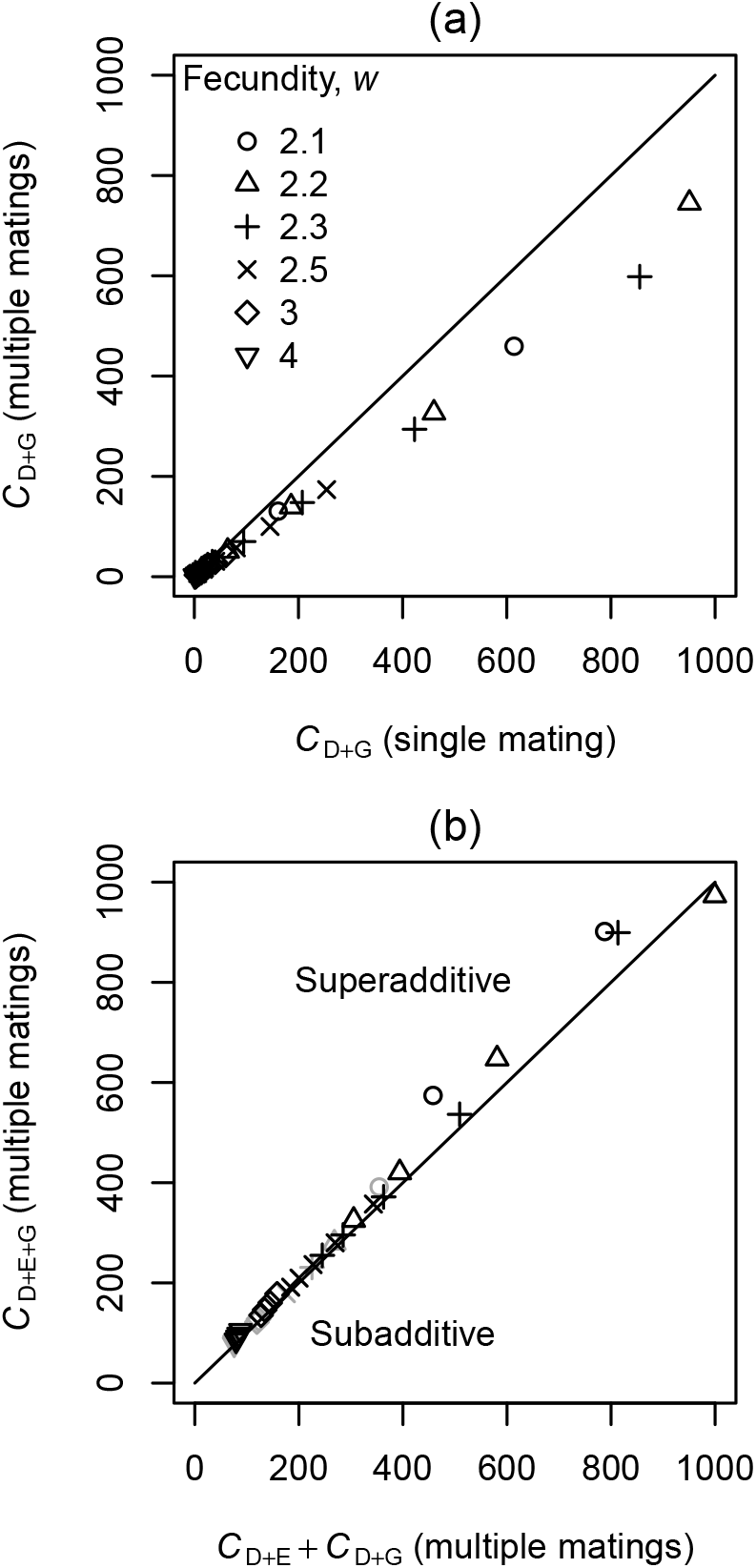
Summary of the results for the model variant with multiple mating events per female. (a) Critical population sizes in the **D+G** scenario with multiple mating events, compared to the case of a single mating event presented in the main text. (b) Relationship between the critical population size under both ecological and genetic problems, *C_D+E+G_*, and the sum of the critical sizes with only ecological problems acting, *C_D+E_*, or only genetic problems acting, *C_D+G_*. Parameter combinations that exhibit a “meaningful” strong Allee effect (success probability at the critical size larger than 1/3) are shown in black, other parameter combinations in gray. In both (a) and (b), the various points correspond to different values of the fecundity, *w*, and the number of lethal equivalents, *b* (those represented in Fig. 2). The diagonal lines represent points where both critical population sizes are equal (a) or the Allee effects are additive (b). Other parameters: *n* = 1,000, *r* = 0.02.

## Appendix S2 Hermaphroditic model version

In this model version, there are no separate sexes but each individual can function both as a male and as a female. We assume, however, that in each mating event, an individuals either acts as a male or as a female, but not both at the same time. Each individual in its role as female finds a mating partner with probability 1 − *e*^−*r*/2·*N_t_*^. If successful, the individual produces a Poisson-distributed number of offspring with parameter *w*/2. The individual acting as father is then chosen at random from the *N_t_* individuals. The parameters *r* and *w* are divided by two to make the results comparable to the two-sex model. The genetic part of the model is exactly as described in the main text.

Critical population sizes with just demographic stochasticity (**D**) were all below one in the hermaphroditic model version, substantially lower than in the model with two sexes where they took values between 2 and 3. This indicates that the slight distortion of success probability curves in the **D**-scenario of the original model was caused by sex-ratio fluctuations in very small populations.

For the other three scenarios, the results were qualitatively very similar to those of the model version with two sexes (Fig. S9). Most importantly, parameter combinations that led to a meaningful strong Allee effect with a relatively small critical population size in the two-sex model were still classified as meaningful. This indicates that the slight S-shape of the corresponding success-probability curves was indeed caused by inbreeding depression and not just by sex-ratio fluctuations. Overall, critical population sizes in the **D+G**-scenario were slightly lower in the hermaphroditic compared to the two-sex model (Fig. S9 a), but the additivity of Allee effects at 1,000 loci was preserved (Fig. S9 b).

**Figure S9:**
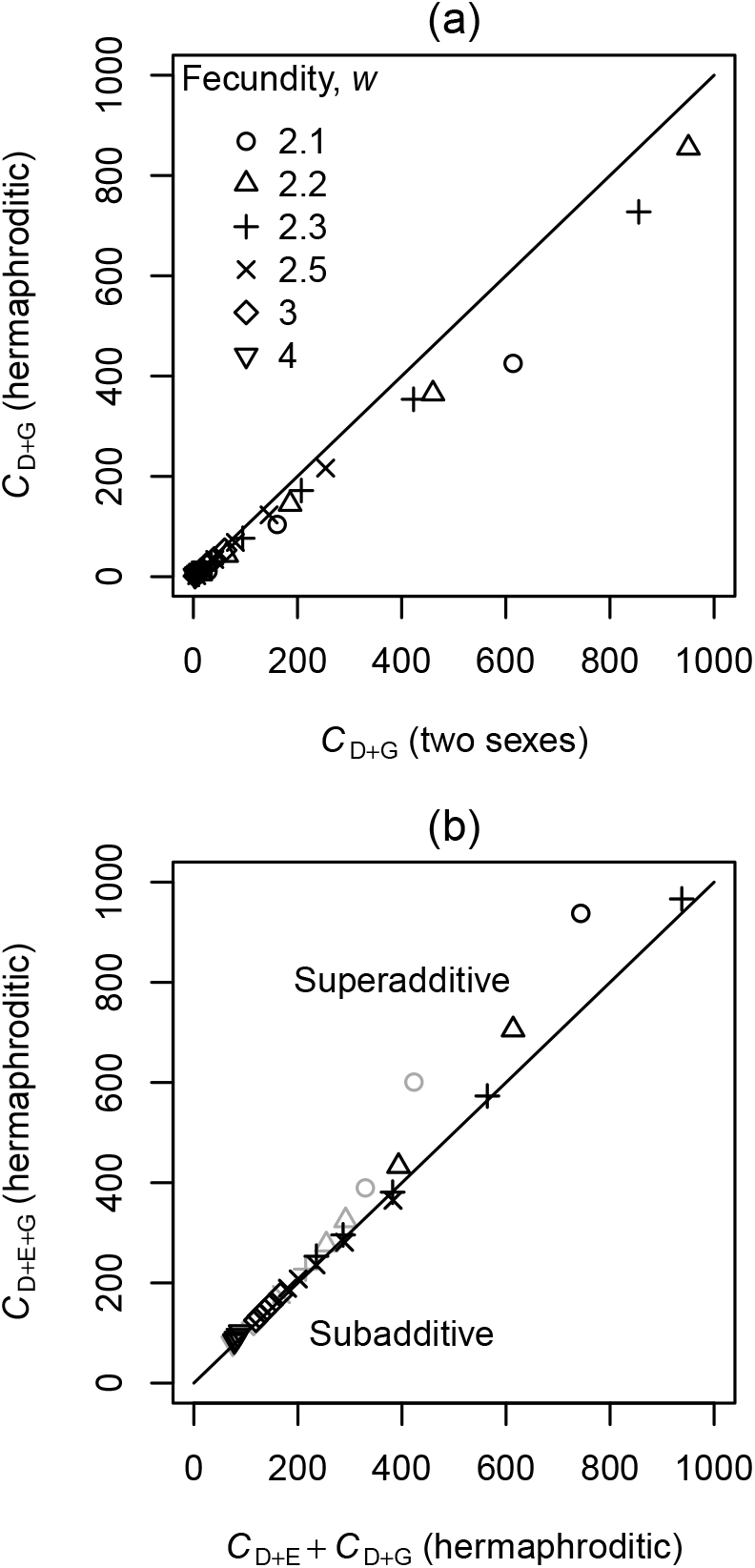
Summary of the results for the hermaphroditic model variant. (a) Critical population sizes in the **D+G** under hermaphroditic mating, compared to the two-sex model presented in the main text. (b) Relationship between the critical population size under both ecological and genetic problems, *C_D+E+G_*, and the sum of the critical sizes with only ecological problems acting, *C_D+E_*, or only genetic problems acting, *C_D+G_*. Parameter combinations that exhibit a “meaningful” strong Allee effect (success probability at the critical size larger than 1/3) are shown in black, other parameter combinations in gray. In both (a) and (b), the various points correspond to different values of the fecundity, *w*, and the number of lethal equivalents, *b*. The diagonal lines represent points where both critical population sizes are equal (a) or the Allee effects are additive (b). Other parameters: *n* = 1, 000, *r* = 0.02.

## Appendix S3 Extended model with alternative genetic architectures

In the main text, we have assumed that all mutations are lethal, fully recessive, and have the same lethal allele frequency in the source population from which the founder individuals are drawn. In this extended model, we introduce a set of additional parameters for each locus *i*, the selection coefficient, *s_i_*, and the dominance coefficient, *h_i_*. In addition, the loci may differ in their frequencies in the source population, *q_i_*. Assuming multiplicative viability selection, the viability, *v*, of an individual with *d_i_* deleterious allele copies at locus *i* is

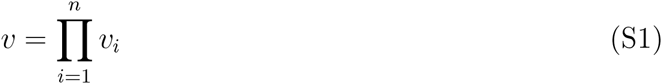

with

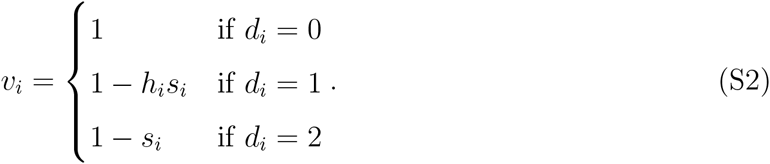

Both *s_i_* and *h_i_* take values between 0 and 1. With *s_i_* = 1 and *h_i_* = 0 for all *i*, we recover the model with fully recessive lethal mutations that is discussed in the main text. To determine whether an offspring individual is viable or not in the modified model, we draw a Bernoulli random variable. With probability *v*, the individual survives to adulthood, and with probability 1 − *v* it dies. As for the original model, we assume that all founder individuals are drawn from the viable members of the source population. We therefore randomly generate founder individuals and perform a Bernoulli draw for their viability until we have the pre-specified number of founder individuals, *N*_0_. Mate-finding and reproduction proceed in the same way as in the original model.

For a given number of lethal equivalents, many different choices are possible for the underlying genetic architecture, i.e. for the number of loci and their individual parameters *s_i_, h_i_*, and *q_i_*. In the following subsections, we will first explain how to compute the contribution of a locus with parameters *s_i_, h_i_*, and *q_i_* to the total number of lethal equivalents, *b*. Then we consider three example genetic architectures to determine whether genetic Allee effects also occur for more complex and possibly more realistic architectures than considered in the main text.

### Appendix S3.1 Lethal equivalents contributed by an individual locus

The standard model for the cost of inbreeding, originally due to Morton *et al*. (1956), is of the form

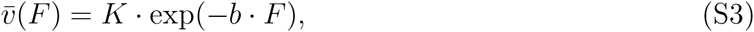

where 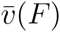 is the average viability for a given inbreeding coefficient *F, K* is a constant summarizing sources of mortality not related to inbreeding, and the parameter *b* is called the number of “lethal equivalents”. To understand the interpretation of *b* suggested by its name, consider a population in which individuals carry on average *b* fully recessive lethal mutations per haploid genome. We further assume that there are many loci such that each mutation is individually very rare in the population. An individual dies from inbreeding if it carries at least one lethal mutation in the portion of the genome *F* that is identical by descent and therefore homozygous. Assuming independence across loci, the number of mutations in this genomic region is approximately Poisson-distributed with mean *b* · *F*, and the probability that there are no mutations in this region is exp(−*b* · *F*). Thus, a value of *b* obtained by fitting (S3) to empirical viability data can be interpreted as an effective number of recessive lethal mutations, i.e. the average number of fully recessive lethal mutations that would produce the same cost of inbreeding as the actual genetic architecture under consideration.

Using (S3) and assuming multiplicative selection and independence across loci, the total number of lethal equivalents, *b*, can be split into contributions of individual loci:

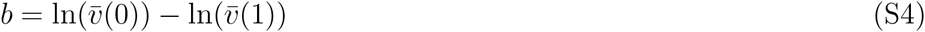

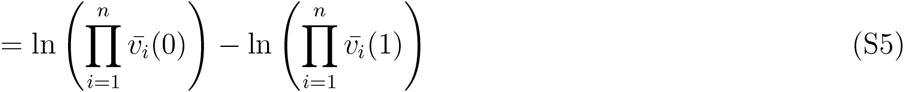

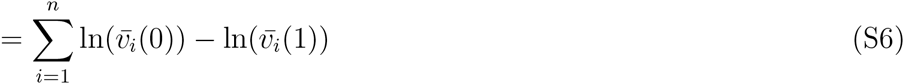

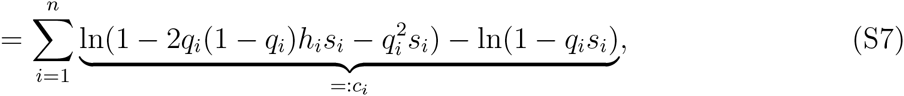

where *c_i_* is the contribution of locus *i*.

Note that if both *q_i_* and *h_i_* are small

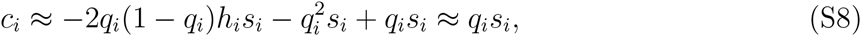

where we used ln(1 − *x*) ≈ −*x* for small *x*.

**Figure S10:**
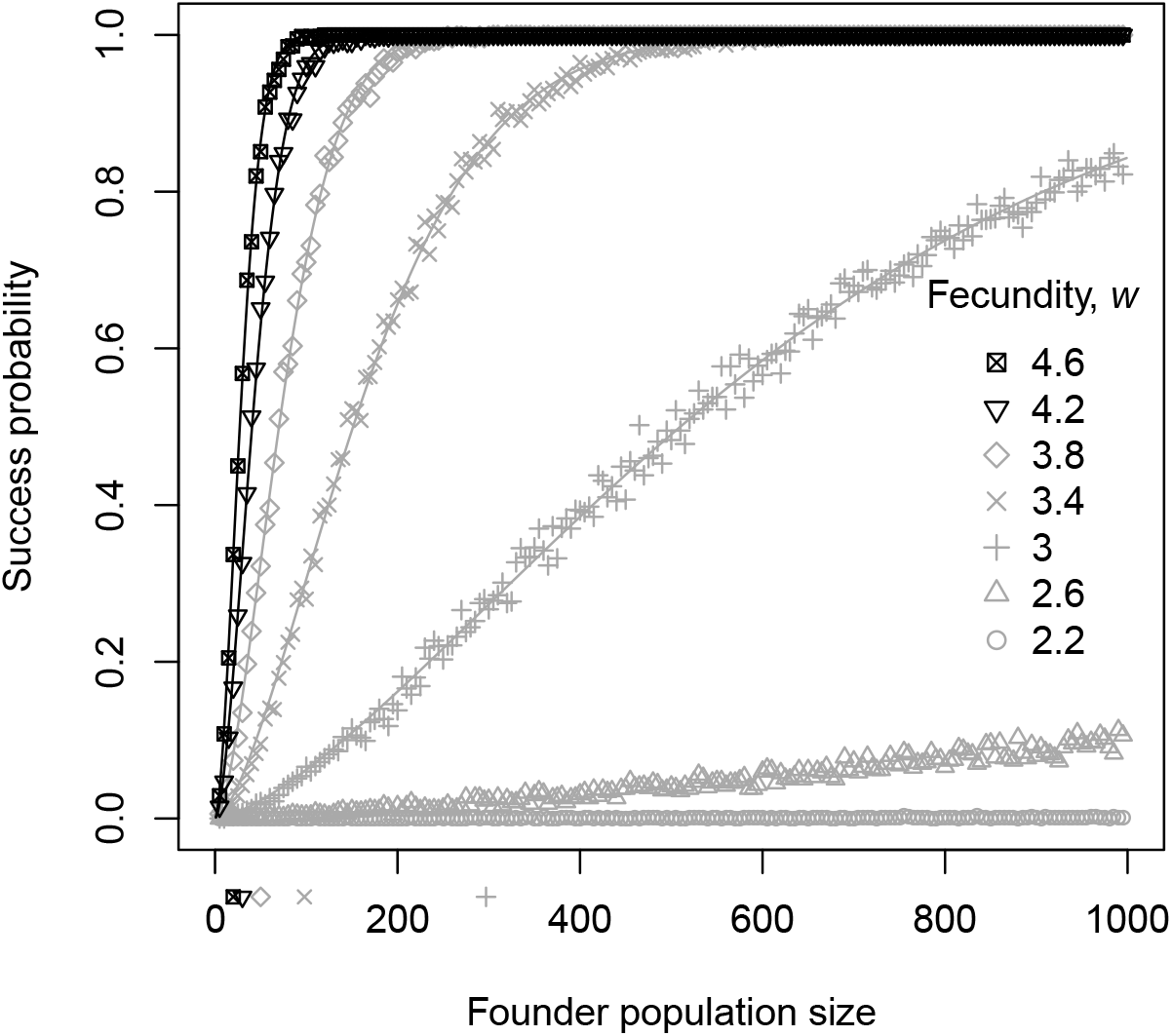
Success probability curves for populations with initially eight lethal equivalents due to lethal mutations with a dominance coefficient *h* = 0.05. Symbols represent success probabilities estimated from the simulations and lines are the corresponding fits of the Weibull model (1). Symbols below the *x*-axes indicate critical population sizes estimated from the Weibull model. Parameter combinations that exhibit a “meaningful” strong Allee effect (success probability at the critical size larger than 1/3) are shown in black, other parameter combinations in gray.

### Appendix S3.2 Lethal mutations with dominance coefficient *h* = 0.05

In this set of simulations, parameters were identical across loci. As in the main text, we ran simulations with different values for the number of lethal equivalents, *b*, and the underlying number of loci, *n*. Assuming equal contributions across loci and using the approximation (S8), we set the allele frequency of lethal mutations in the source population to *q* = *b*/*n*.

With this choice of parameters, success probabilities were overall lower than with *h* = 0, which is expected since the mutations now have an additional deleterious effect in heterozygotes which reduces overall population viability. More interestingly, also the shapes of the success probability curves changed. The S-shape was less pronounced and few parameter combinations were classified as having meaningful genetic Allee effects. Fig. S10 shows examples for success probability curves with *b* = 8 and for various fecundity values, *w*.

### Appendix S3.3 Joint distribution of selection and dominance coefficients estimated by Agrawal & Whitlock (2011) for yeast

Agrawal & Whitlock (2011) used growth-rate data from thousands of knockout mutants in the yeast *Saccharomyces cerevisiae* and a maximum-likelihood modeling approach to estimate the joint distribution of selection and dominance effects for mutations in yeast. We use their best-fitting model (Model 7) and the associated parameter estimates. However, we only model deleterious mutations. Specifically, for 23.5 % of mutations, the deleterious selection coefficient was drawn from a uniform distribution on the interval [0, 0.304]. The remaining mutations had their selection coefficient drawn from an exponential distribution with mean 0.009 in 88.6 % of the cases and from an exponential distribution with mean 0.04 in the remaining cases. Selection coefficients larger than 1 were set to 1, i.e. lethality. Conditioned on the selection coefficient, *s*, for a locus, its dominance coefficient was drawn from a Gamma distribution with shape parameter 0.14 and scale parameter 0.26, shifted such that the mean dominance coefficient was 4.23/(1 + 297.4 · *s*). We restricted dominance coefficients to the interval [0,1]. Although this empirical distribution is the most detailed information on the joint distribution of selection and dominance coefficients existing for any organism to date, note that it comes with a number of assumptions and empirical constraints (see detailed discussion by Agrawal & Whitlock 2011). For example, many strong-effect mutations were probably missing from the data set because their growth rates could not be estimated.

A mutation’s selection coefficient and dominance coefficient will clearly influence its frequency in the source population, but genetic drift also plays a role. For a locus *i* with selection coefficient *s_i_* and dominance coefficient *h_i_*, we therefore drew an allele frequency *q_i_* from Wright’s distribution for allele frequencies at mutation-selection-drift balance (Wright 1937). Specifically, given a source population size, *N_s_*, and a lethal mutation rate, *μ*, and ignoring back-mutation, the probability density of lethal allele frequency, *q_i_*, is approximately

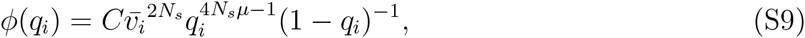

where *C* is a normalization constant and 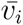 is the mean viability at locus *i*, i.e. 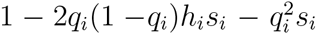. Some example probability densities for the allele frequency of fully recessive lethals are shown in Fig. S11. In the following, we assume a source population size *N_s_* = 10, 000 and a per-locus mutation rate of *μ* = 10^−6^.

To build a genetic architecture underlying a given number of lethal equivalents in the source population, we proceeded as follows. As long as the cumulative number of lethal equivalents was below the desired value, we generated a new locus with parameters *s_i_, h_i_*, and *q_i_* using the model outlined above, then used (S7) to compute its contribution, *c_i_*, and added the result to the current number of lethal equivalents. Using this method, we would have needed about 60,000 loci per lethal equivalent, which would have required too much time and memory to simulate. We therefore sorted the mutations according to their contributions, *c_i_*, to the lethal equivalents. Since a portion of very dominant mutations contributed negatively to the lethal equivalents, we could still obtain the desired number of lethal equivalents by taking a smaller number of the strongest contributors. For example, for 2 lethal equivalents, we needed about 6,000 mutations. Since the presence of these 6,000 mutations substantially reduced the average viability in the population, we had to choose higher values of the fecundity, *w*, than in the main text to avoid the extinction of all populations. Example results are shown in Fig. S12.

**Figure S11:**
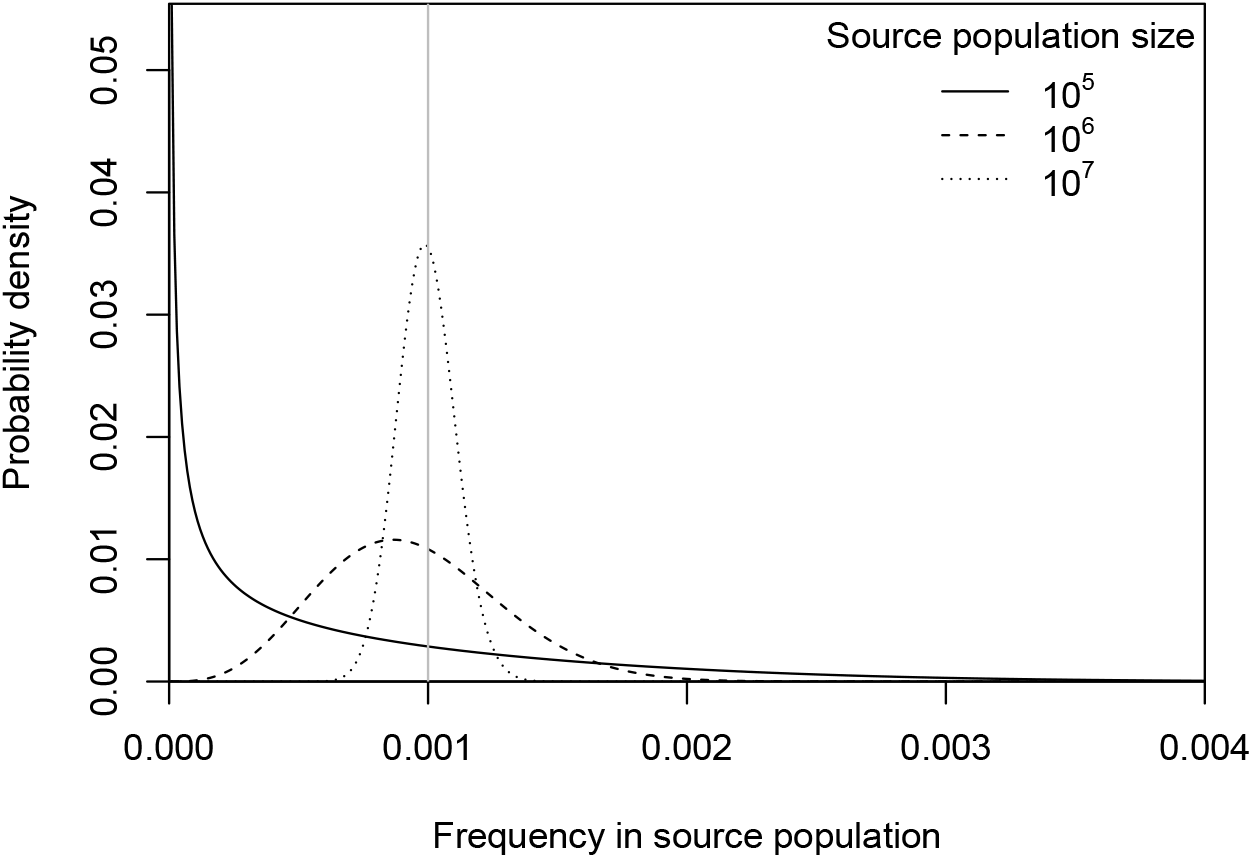
Probability density functions for lethal mutant allele frequencies in source populations of different sizes. The vertical gray line indicates, 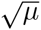, the equilibrium frequency under mutation-selection balance in an infinitely large population. *μ* = 10^−6^.

**Figure S12:**
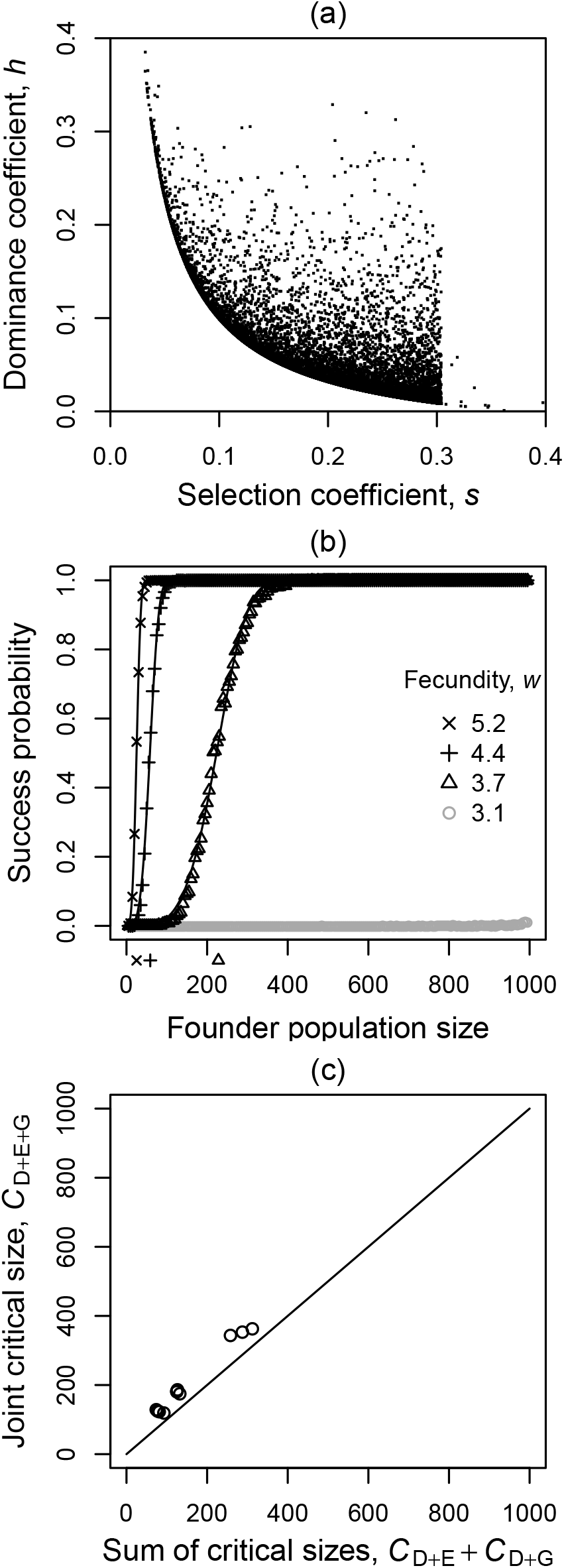
Results with parameters of individual loci sampled from the joint distribution of selection and dominance effects estimated by Agrawal & Whitlock (2011). (a) Selection and dominance coefficients for the 17,828 contributing loci underlying six lethal equivalents. (b) Success probability curves with six lethal equivalents and various values of the fecundity, *w*. Symbols represent success probabilities estimated from the simulations and lines are the corresponding fits of the Weibull model (1). Symbols below the *x*-axes indicate critical population sizes estimated from the Weibull model. For *w* = 3.1, success probabilities were overall too low to estimate a critical population size. (c) Relationship between the critical population size under both ecological and genetic problems, *C_D+E+G_*, and the sum of the critical sizes with only ecological problems acting, *C_D+E_*, or only genetic problems acting, *C_D+G_*. The points represent different values of *b* and *w*. The Allee effect was meaningful in all cases shown, i.e. for all parameter combinations for which the model could be fit.

### Half lethals, half small-effect mutations

Here, we considered the extreme case where all mutations are fully recessive (*h_i_* = 0 for all *i*). Half of them were lethal and the other half had their selection coefficient drawn from an exponential distribution with mean 0.05. Selection coefficients larger than one were set to one. Wright’s distribution (S9) with 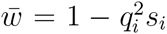 was used to draw allele frequencies for the individual mutations in the source population. For lethal mutations and mutations of smaller effect separately, the lethal equivalents contributed by randomly generated mutations (S7) were then summed up until the desired value was reached. In this case, the resulting number of loci was small enough to be used directly in the simulation and no viability adjustment was made. Representative success-probability curves and results on the interaction of Allee effects are shown in Fig. S13.

**Figure S13:**
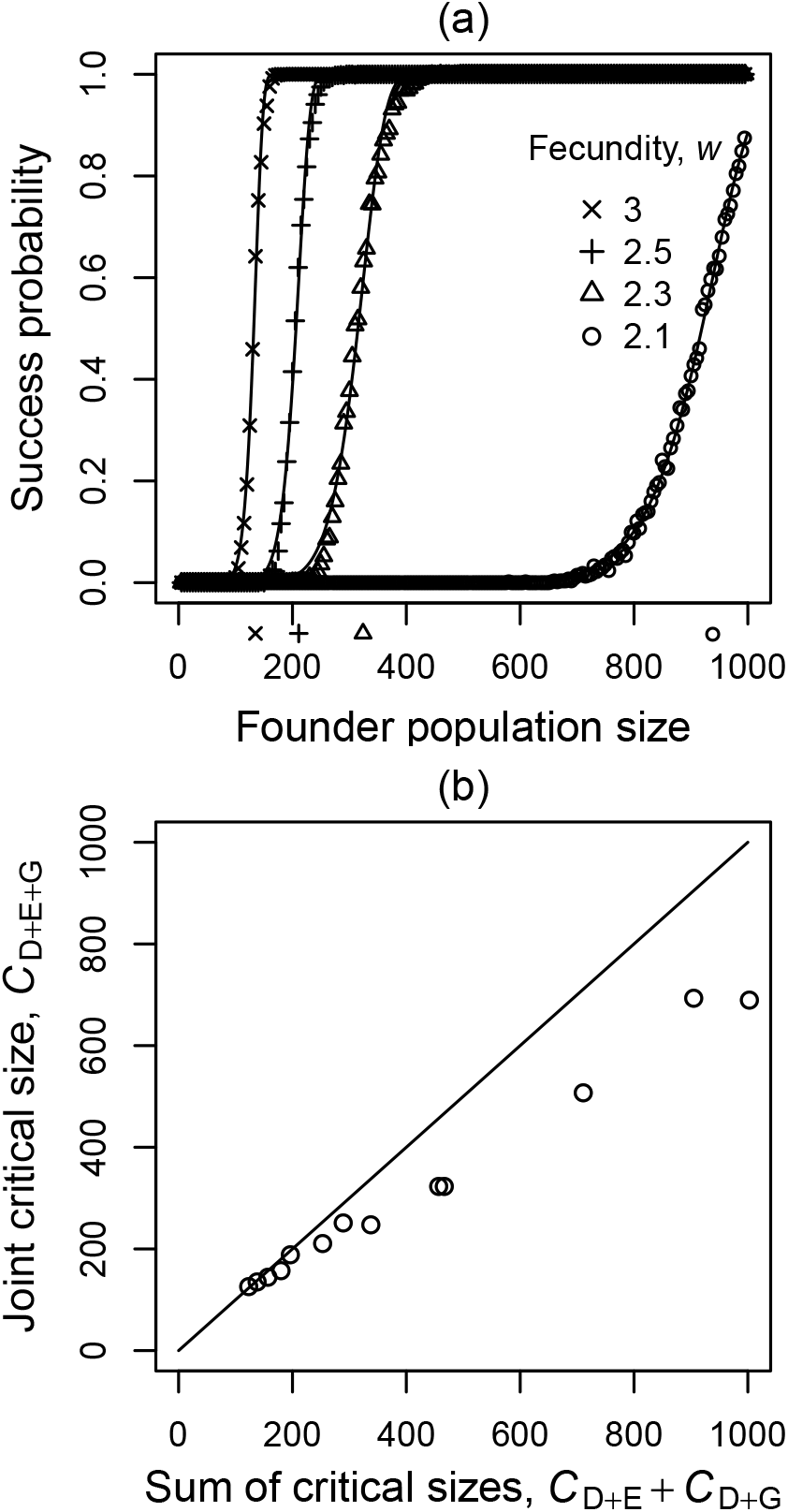
Results for a genetic architecture with half of the lethal equivalents due to fully recessive lethals and the other half due to fully recessive mutations of smaller effect. (a) Success probability curves for populations with initially four lethal equivalents. Symbols represent success probabilities estimated from the simulations and lines are the corresponding fits of the Weibull model (1). Symbols below the x-axes indicate critical population sizes estimated from the Weibull model. In this case, all parameter combinations exhibit a “meaningful” strong Allee effect (success probability at the critical size larger than 1/3). (b) Relationship between the critical population size under both ecological and genetic problems, *C_D+E+G_*, and the sum of the critical sizes with only ecological problems acting, *C_D+E_*, or only genetic problems acting, *C_D+G_*. The points represent different values of *b* and *w*. For this set of simulations, all genetic Allee effects were meaningful in the sense that the success probability at the critical size was larger than 1/3.

## Appendix S4 Analytical results for single-locus model

In this section, we derive approximate expressions for the survival probability *P*(*N*_0_) of a small founder population of size *N*_0_ in the absence of mate limitation and with lethal, fully recessive alleles occurring at a single locus, and we show that *P*(*N*_0_) does not exhibit the characteristic signature of a strong Allee effect.

To start deriving *P*(*N*_0_), it is important to notice two key points. (1) The population survives if and only if the wildtype, i.e. healthy, allele does not go extinct. This is easy to see because mutant homozygotes are not viable. (2) Since with fully recessive mutations heterozygotes and wildtype homozygotes have the same fitness, the number of descendants in the next generation of the various wildtype allele copies in the population are identically distributed. Thus, the distribution of the number of descendants stays constant and, in particular, does not depend on the total population size or the number of wildtype and mutant alleles. The numbers of descendants of the various wildtype allele copies are not completely independent though. For example, if the population consists just of one female wildtype homozygote and one male wildtype homozygote, the allele copies sitting in the male can only reproduce if those sitting in the female reproduce as well. However, these dependencies should be weak, especially in slightly larger populations. Also this dependence is unrelated to genetic structure and it is present even with just demographic stochasticity. Therefore, we will approximate the dynamics of wildtype allele copies in the population by a homogeneous branching process.

Given an initial copy number of the wildtype allele, *Z*, the probability that the population survives is 1 − *u^Z^*, where *u* is the probability that a single wildtype lineage goes extinct. In our model, *u* is a function of the average number of offspring per mated pair, *w*_0_, but for the present purpose it only matters that it does not depend on *Z* or *N*_0_.

To derive *P*(*N*_0_), we have to average over all possible configurations of the founder population:

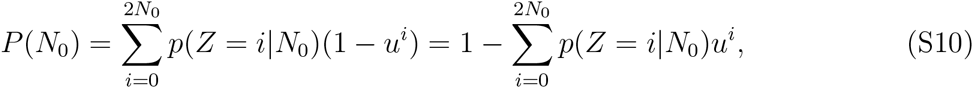

where *p*(*Z* = *i*|*N*_0_) is the probability that a founder population with *N*_0_ individuals contains *i* copies of the wildtype allele.

In our simulation model, we have assumed that the founder individuals are drawn from the viable members of the source population, which means that a founder individual is a wildtype homozygote with probability (1 − *q*)^2^/(1 − *q*^2^) and a heterozygote with probability 2*q*(1 − *q*)/(1 − *q*^2^). Hence *N*_0_ ≤ *Z* ≤ 2*N*_0_ and with *Y* = 2*N*_0_ − *Z* as the number of heterozygotes (equal to the number of mutant alleles) in the founder population

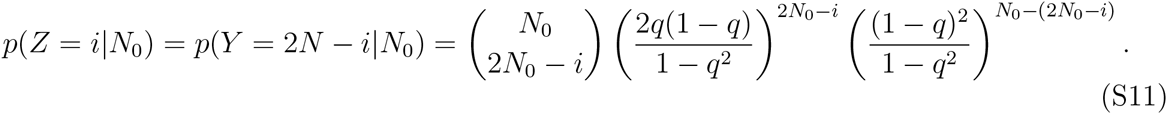

With this,

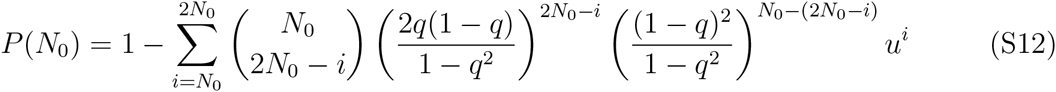

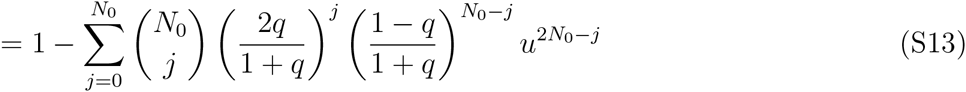

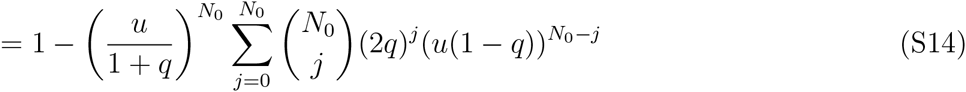

Using the binomial theorem, we finally obtain

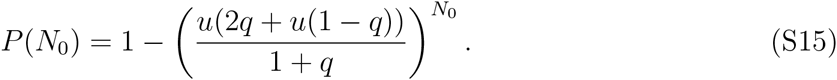

Thus *P*(*N*_0_) is of the form

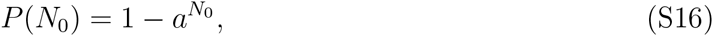

where *a* is a positive expression that depends on *q* and potentially also on other model parameters via *u* but, importantly, does not depend on *N*_0_.

To check for the presence of a strong Allee effect, we consider the second derivative of *P*(*N*_0_),

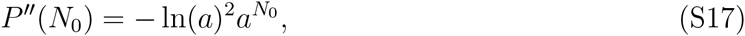

which is always negative. Therefore, the success probability curve does not have an inflection point and there is no strong Allee effect. In fact, the shape of the success probability curve is the same as for a model with just demographic stochasticity, which also can be represented as a homogeneous branching process.

The same holds true for some other assumptions on how the founder population is constructed. For example, if we assume that the number of wildtype alleles in the founder population is deterministic and exactly proportional to *N*_0_, i.e. *Z* = *bN*_0_, then

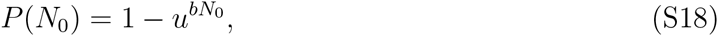

which is again of the form (S16) with *a* = *u^b^*. As another example, if the founder individuals are drawn from the source population before viability selection acts, *Z* is binomially distributed with parameters *q* and 2*N*_0_. Thus

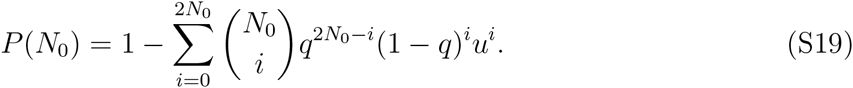

Using the binomial theorem again,

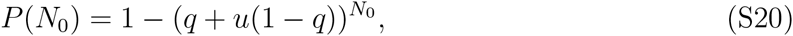

which is again of the form (S16) and thus does not exhibit a strong Allee effect.

